# Self-severing circuits facilitate passage of ingestible electronic sensor-guided therapeutics

**DOI:** 10.64898/2026.03.27.714561

**Authors:** Sean Healy, Angsagan Abdigazy, McKenna Clinch, Jason Y Chin, Mohammad Shafiqul Islam, Zachary Lee, Julia Z Ding, Joy Jackson, Ramy Ghanim, Xavier Manigault, Sritej Ponna, Megan C Lee, Jihoon Park, Yasser Khan, Alex Abramson

## Abstract

Ingestible electronics enable the tracking and treatment of gastrointestinal and systemic diseases. However, bulky batteries and circuit boards require large capsules that can result in bowel obstruction, a medical emergency. Here, we engineered a 9 × 26 mm electronic pill capable of triggered severing into tiny pieces with sizes clinically proven to reduce obstruction risk. Our capsule enables multicomponent circuit boards to connect with separately encapsulated powering elements via conductive, interlocking connections. Heat induced softening of polyethylene glycol/polycaprolactone channels activates a spring to separate encapsulated components into inert 9 × 15 mm segments, facilitating intestinal passage. Separation triggers included closed-loop sensors and time-delay circuits. In vivo swine studies demonstrate the ability of our capsules to sense luminal oxygen changes via an optoelectronic sensor, locally trigger upadacitinib delivery, and facilitate safe excretion.

## Introduction

Electronic pills bring sensors and actuators next to gastrointestinal tissues, luminal fluids, and vasculatures that enable unique diagnostic and therapeutic interventions^1–3^. They empower healthcare professionals to remotely execute minimally invasive endoscopies^4^, track patient adherence to medications^5^, perform enteric stimulation therapies^6^, as well as sense physical properties and chemical analytes within the gastrointestinal (GI) tract^7^. Preclinical devices also support extended or triggerable drug delivery^8,9^ and electrical stimulation therapy^10^. Furthermore, patients widely prefer pills as a means of diagnostic or therapeutic intervention over patches, implants, and injections^11,12^.

As the complexity of ingestible electronics expands to support a wider range of diagnostic and therapeutic functions, their increasing size limits their safety profile^13^. Electronic pills require large, nondegradable powering, communication, and actuation components that increase capsule size and risk generating bowel obstructions. A bowel obstruction is a medical emergency that can develop rapidly when objects partially or completely block fluid, food, and gas from passage^14^. Due to the risk of bowel strangulation and perforation, suspected obstruction requires immediate imaging and surgery consultation to prevent necrosis, perforation, infection, and sepsis^15^. Therefore, healthcare professionals must constantly monitor their patients when administering electronic capsules until passage occurs. This greatly limits the applicability of ingestible electronics to single use rather than daily or weekly dosing, which is required for applications like drug delivery or continuous sensing.

Capsule size significantly impacts rates of gastrointestinal obstruction. Daily dosed, nondegradable osmotic pump capsules measuring 9 × 15 mm demonstrated 0 reported gastrointestinal obstruction cases in 22 million applications^16^. Comparatively, the 11 × 26 mm PillCam, dosed once under professional supervision, generated gastrointestinal obstructions in 13% of patients with known Crohn’s diseases^17^ and in 1.4% of patients with suspected bowel disease^18^. FDA cleared and approved ingestible electronics utilize commercial components that inhibit capsules from reaching 9 × 15 mm dimensions. Inspired by a lizard’s ability to detach its tail to escape predation^19^, here we describe a modular electronic capsule capable of autonomous disassembly into pieces ≤ 9 × 15 mm after 24 hours in the body.

In this paper we introduce the ESCAPE platform, Excretable Self-Cleaving Autonomous Pill Electronics. ESCAPE allows nondegradable circuit boards and powering elements to be ingested as a single, watertight capsule and then sever into passable components on command to facilitate GI passage (**Fig 1A**). To electrically join capsule segments, we engineered a universal, press-fit mechanism with < 0.5 Ω of resistance. To separate the segments at their interface, we fabricated a 1.43 mWh, heat-triggered actuation mechanism that releases a compressed spring to propel segments apart. Triggering occurs via guidance by an onboard sensor, manually through Bluetooth, or automatically after a set period in the body to reduce obstruction risk. We also demonstrate that ESCAPE enables sensor triggerable, localized drug delivery of up to 20 mg payloads following device separation. Here, we utilized our capsule’s unique severing capability to trigger drug delivery at specific oxygen thresholds that mimic those found in the inflamed intestinal lumens of patients suffering from Crohn’s Disease or Ulcerative Colitis.

**Figure 1:**
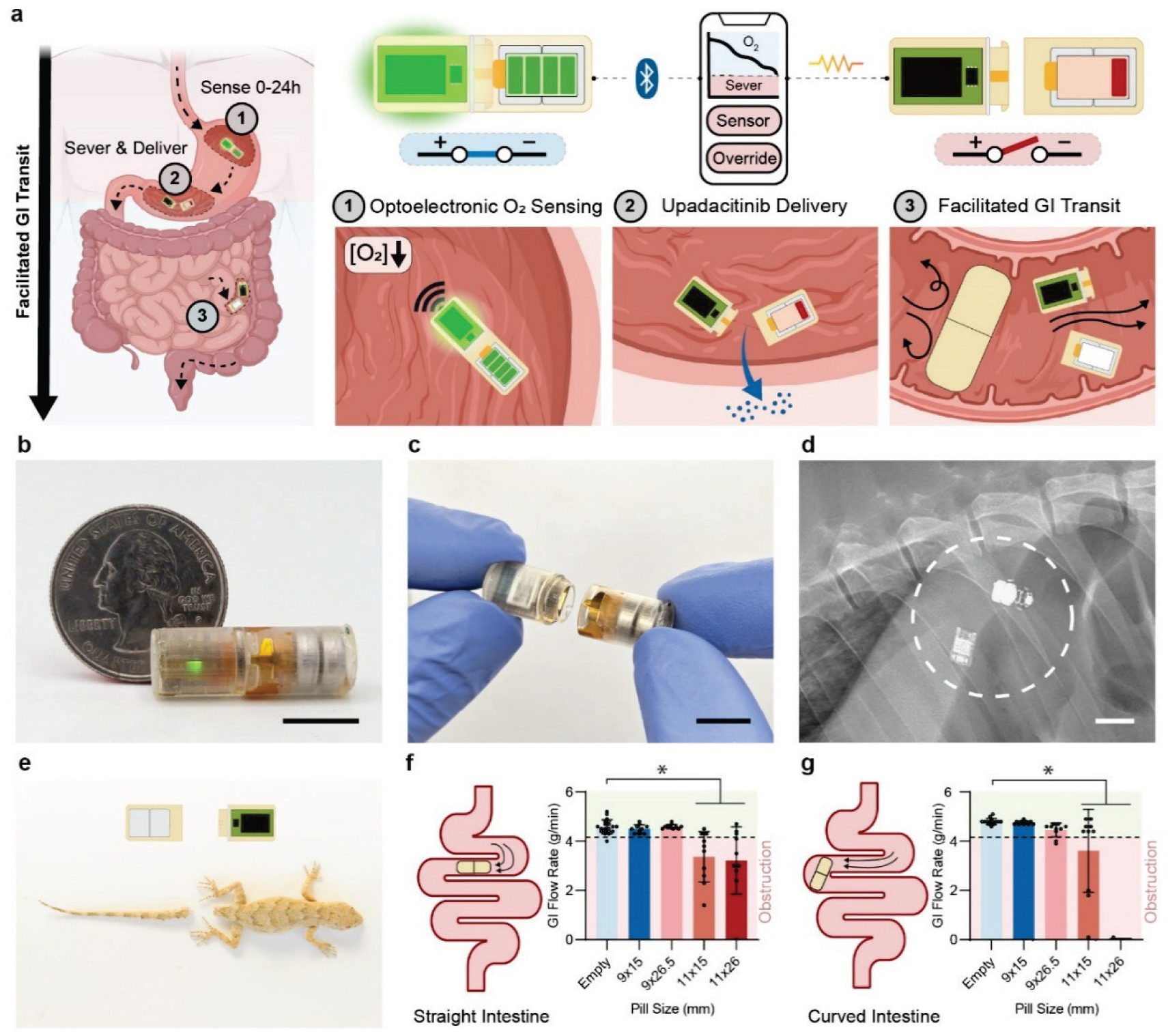
**a,** The 9 × 26 mm ESCAPE capsules facilitate GI passage of large, non-degradable electronic components by autonomously severing into two 9 × 15 mm segments. Mechanical and electronic interlocks are severed via either manual input or autonomously via on-board electronics. Example ESCAPE activities include optoelectronic gas sensing and closed-loop drug delivery. **b,** Joined and **c**, severed ESCAPE platforms. **d,** Radiograph of ESCAPE following severing in an in-vivo swine model. **e,** ESCAPE’s severing mechanism is inspired by autotomy, a biological mechanism employed by animals such as lizards to voluntarily self-amputate a pre-defined limb. **f,** Ex-vivo swine models of straight and curved intestinal junctions test for intestinal obstructions utilizing flowing chyme, with obstruction defined as a < 4.1 g/min flow rate. Capsule dimensions of 9 × 15 mm were the only size to not generate obstruction in either geometry (Avg ± SD, dots = individual trials. n = 10 devices, n = 15 - 20 empty trials. straight SI, p = 0.0001, p = 0.0001; curved intestine, p =, 0.0003, 0.0001; ANOVA, one-tailed Dunnett’s test). Illustrations in **a** created with Biorender.com and **a**, **e** licensed from Adobe Stock under Adobe For Enterprise License. Scale bars = 10 mm.

In vivo in swine, we demonstrate that ESCAPE enables continuous optoelectronic monitoring of surrounding luminal oxygen levels in addition to the locally triggered delivery and uptake of upadacitinib^20^. One of the only consistent indicators for gastrointestinal organ targeting is luminal oxygen concentration, which decreases from 8-15% in the stomach down to 2-4% in the small intestine and 0-1% in the colon^21–23^. Oxygen levels also act as an indicator for multiple GI disorders, including inflammation present in Crohn’s Disease and Ulcerative Colitis^22–25^ Here, we controlled oxygen levels in the GI tract by insufflating air and CO2 in the stomach to demonstrate that ESCAPE can sense physiological changes in luminal oxygen levels. We then triggered the severing of ESCAPE to deliver upadacitinib, an oral Janus Kinase inhibitor that is FDA-approved for the treatment of inflammatory bowel disease but also generates severe systemic side effects when delivered in its current formulation^26^. Additionally, we showed through computational, ex vivo, and in vivo swine models that ESCAPE facilitates passage through the GI tract while enabling the ingestion of large electronic components.

### Device Design for Facilitated GI Transit

The ESCAPE platform severs itself from a 9 × 26 mm combined form factor into two 9 × 15 mm segments to facilitate passage through the GI tract (**Fig 1B-1D**). This severing capability is inspired by autotomy, the capability of certain animals, such as lizards, to voluntarily self-amputate pre-defined body parts to escape predation (**Fig 1E**). ESCAPE implements a similar structure where form factors connect to each other using simple geometries such as grooves and ridges. Each segment casing is compatible with injection molding techniques and materials to facilitate future clinical translation. Capsule electronics exposed to the GI tract environment after severing are either fabricated from known biocompatible materials or masked with medical grade UV-epoxy and parylene-C.

We selected capsule segment dimensions of 9 × 15 mm to minimize the risk of GI obstruction as the severed ESCAPE platform passes through the GI tract. To test the obstruction risk of different capsule dimensions, an ex vivo swine intestine set-up was built to mimic the flow of chyme through the straight and curved pathways of the healthy intestinal environment. Using this experimental set-up, we confirmed that 9 × 15 mm capsules do not cause obstruction in either a straight or curved intestinal path (**Fig 1F, 1G**). We also demonstrated that 11 × 15 mm and 11 × 26 mm capsules generated obstructions in our intestinal model. These results are similar to the obstruction risk seen with similarly sized osmotic pump capsules and PillCams in humans, respectively. We also tested a 9 × 26 mm capsule, which did not generate obstruction in the straight segment model but can generate obstruction in the curved model. The observed results in straight intestinal pathways agree with COMSOL simulations conducted with the same capsule geometries, establishing symmetry between both theoretical and experimental evaluations of capsule obstruction (***supplementary figure 1***).

### Joinable Electrical Contacts for Capsule Modularity

ESCAPE’s joinable structure links two distinct capsule segments to form stable, low impedance electrical connections that can be severed through mechanical separation (**Fig 2A**). Here, one segment possesses a power source while the other contains a printed circuit board (PCB).

**Figure 2:**
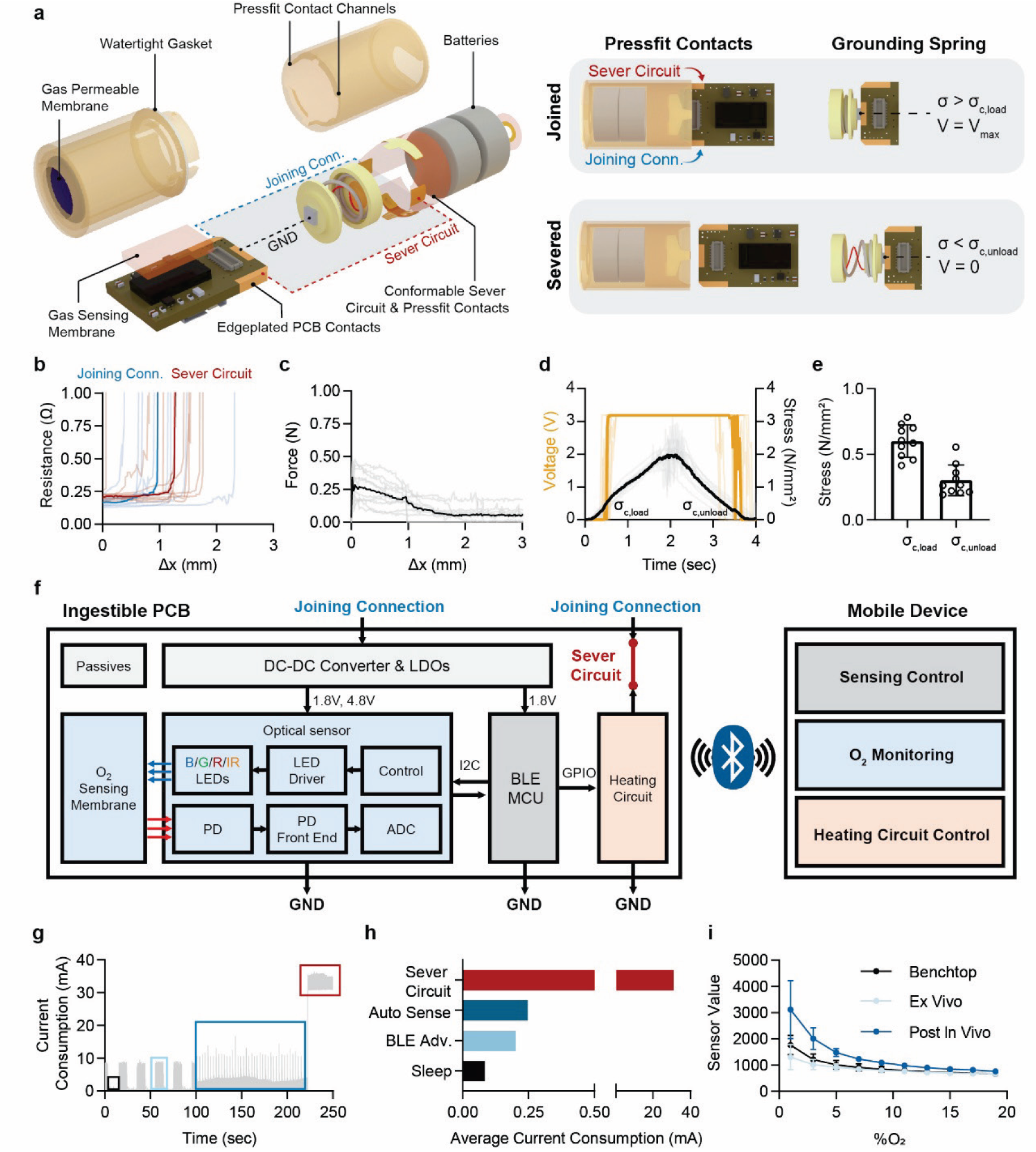
**a,** Exploded view of device components. Three electronic pathways exist between the power and control capsule segments: (1,2) Two pressfit contacts connecting edge-plated rails on the control capsule’s PCB to the power capsule and (3) a pressure sensitive grounding spring that directly connects ground to the central edge-plated contact on the PCB. **b,** The pressfit rail contacts engage to form sub-0.3Ω impedance pathways that are easily disengaged with minimal force (n = 10 devices, bold = median). **c,** The grounding spring switches from high-to-low impedance configurations with sufficient applied stress, where the stress required to go from high-to-low is greater than low-to-high (Avg ± SD, n = 10 device), bold = median). **d**, Block diagram for PCB functions and data transmission. **e**, Device current consumption during different PCB functions. **f**, Average current consumption for each PCB function. **g**, ESCAPE’s optoelectronic sensor retains a high response to decreased oxygen concentration during and after exposure to physiological conditions (Avg ± SD, n = 5 membranes).

Pressfit contacts between segments connect circuit components via a flexible printed circuit board (fPCB) that conformally wraps around the capsule’s inner wall (***supplementary figure 2***). The positive battery terminal is connected to edge-plating on the PCB via a sliding press fit rail. Similarly, a second rail connects side plating on the other side of the PCB to a severing circuit that triggers capsule separation. The severing circuit is described in the next section. These links possess negligible interfacial impedance and allow for minimal frictional resistance during severance (**Fig 2B, 2C**). The contacts are designed to be universally compatible with any 8.1 mm wide rectangular PCB that includes edge-plated sides.

Front edge-plating on the PCB forms the final connection with the negative battery terminal via a pressure-based connection utilizing a compressed spring (**Fig 2D, 2E).** The grounding spring holds a conductive pad in contact with the edge-plating while allowing a wire to connect back to the battery. This spring also provides mechanical force to eject the capsule segments when the severing circuit activates. To prevent unwanted current leakage once severed, the grounding spring is masked with a pressure sensitive quantum tunneling composite (QTC) electrode that naturally exhibits high impedance (>30 MΩ) in an unstressed state. Previously, this material was used to eliminate battery-induced damage when inserted in the GI tract^27^. In a joined state, the inserted PCB overcomes the electrodes activation stress of 0.60 ± 0.13 N/mm^2^, creating a low impedance ground terminal. When severed, the QTC returns to its high impedance state as the applied stress falls to less than 0.30 ± 0.12 N/mm^2^. As a result, ESCAPE can transition between its joined, electrically active state to a severed, electrically inert state that prevents tissue damage during GI passage.

A key advantage of ESCAPE’s joinable structure is its compatibility with interchangeable off-the-shelf electronics that support rapid customization and clinical translation. Compared to smaller custom application-specific integrated circuits (ASICs) that integrate all components into a single chip and require years of development, ESCAPE allows for the implementation of larger commercial components while maintaining a 9 × 15 mm passage size (***supplementary table 1*)**. As a result, changing ESCAPE’s modular functions requires no physical changes to the capsule design.

Our example ingestible PCB (PCB) contains an 8.65 mm × 6.4 mm nRF52 microcontroller (MCU) with Bluetooth Low Energy (BLE) and a 3.5 mm × 7 mm optical sensing module (**Fig 2F, *supplementary figure 3, 4***). The MCU communicates with the optical sensor over the I2C protocol to perform gas sensing measurements. Severing is activated by the MCU through a NMOS transistor switch to trigger current flow through the sever circuit, which is triggered from a mobile device via 2.4 GHz Bluetooth low energy (BLE) communication.

The integration of power-hungry commercial components into ESCAPE’s joinable structure can be achieved without impacting its severing capability. A tradeoff of using commercial components rather than ASICs is their tendency for high current consumption^1^. ESCAPE minimizes these drawbacks by enabling greater volumes for batteries without risking GI obstruction. We utilize two SR41W silver oxide batteries in our capsule, considered as a safe powering option for ingestible electronics^2^. Our PCB further implements sleep cycling of its active electronics to achieve low power operation (**Fig 2G, 2H**). Power management of on-board electronics is especially critical to maximize the batteries’ ability to deliver a final, high current burst for severing. Under an operating lifetime of 24 hours, severing can still be achieved under a constant drain of 0.61 ± 0.01 mA, well below our current consumption rates (***supplementary figure 5***).

As a proof of concept for ESCAPE’s modular electronics, we demonstrated its ability to detect oxygen gas concentrations using an optoelectronic sensor. To detect oxygen gas, our ingestible PCB carries a sensor module previously described in Abdigazy et. al^28^ that utilizes fluorescent quenching for the detection of multiple gases (***supplementary figure 6***). This optoelectronic sensor allows ESCAPE to sense oxygen changes when fully surrounded by ex vivo intestinal tissue, and it remains sensitive after functioning in an in vivo swine stomach (**Fig 2I**).

### Autonomous Severing for GI Transit & Drug Delivery

ESCAPE’s severing functionality is achieved using a compact heat-triggerable ejection mechanism (**Fig 3A, 3B**). In a joined state, capsule segments are interlocked by opposing 0.5 mm tall by 1 mm long trapezoidal channels cast with a blend of polycaprolactone (PCL) and polyethylene glycol (PEG) **(Fig 3C, *supplementary figure 7)***. Inspired by dovetail joinery, these low-profile channels repeatedly withstand 4.5 N of force over 1000 cycles, which is greater than the 2 N forces exerted by the GI tract^29^ plus the 2.5 N working load of the internal grounding spring. However, at temperatures elevated above body homeostasis, the polymer-cast channels soften to provide an unobstructed pathway for capsule severing. Mixing PCL with PEG leads to reduced melting temperatures^30^, and we demonstrate that a 2.5% w/w blend of PEG to PCL negligibly affects channel fracture load (**Fig 3D**).

**Figure 3:**
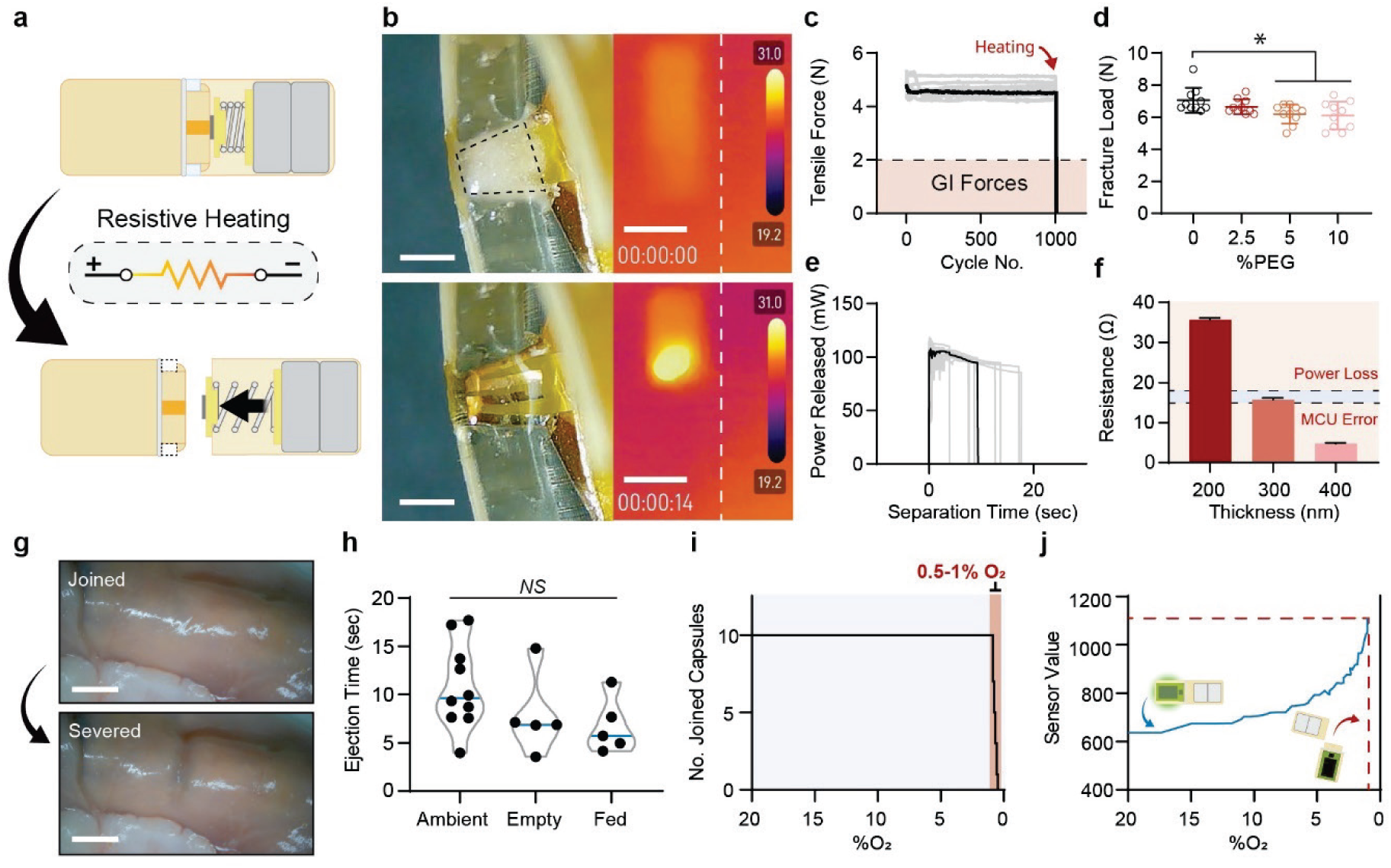
**a,** ESCAPE’s heat triggerable severing and ejection mechanism. **b,** Visual and infrared snapshots of polymer dovetail channels (left scalebar = 1mm; right scalebar = 10 mm). During a sever event, localized hot spots are produced by an inlaid thin film resistive heater, converting the channels from a blocked, load-bearing state to a melted, unobstructed state. **c,** Cyclic durability testing of ESCAPE’s load bearing channels in a solid state, showing that they resist repeated 4.5N loads over 1000 cycles (bold=median, n = 10 devices). **d,** Peak fracture loads for different polymer channel compositions, where the addition of 2.5% melt-mixed PEG negligibly impacts peak fracture load (Avg ± SD, n = 10 devices; p-value =0.0422, 0.0193, ANOVA with Tukey’s). **e,** Power dissipated across ESCAPE’s thin-film resistive heaters when loaded with pre-drained batteries (bold=median, n = 10 devices). **f,** Resistance of thin film resistive heaters as a function of heating layer thickness, where optimal coil resistance ranges from 15 - 18 Ω (Avg ± SD, n = 16 devices). **g,** Ex vivo swine intestine sever tests conducted in empty and fed states showed no statistically significant difference in ejection times (Avg ± Range, n = 5-10 devices, scalebar = 5 mm). **h,** Closed-loop demonstration of ESCAPE’s severing capability, where capsules severed at a pre-programmed O2 threshold of 0.5-1% (n = 10 devices). **i,** Representative curve of ESCAPE closed-loop oxygen sensing and severing at a prescribed threshold. Illustrations in **a** created with Biorender.com and images licensed from Adobe Stock under Adobe For Enterprise License.

To trigger a severing event via on-command polymer melting, current is passed through a conformable resistive heater built into each polymer channel base. Our silver oxide batteries supply enough power to achieve severing within 20 seconds after 24 hours of constant 0.6 mA drain, a 2x worst-case power consumption for any on-board electronics (**Fig 3E**). To convert this applied power into localized heating, we designed thin-film (12.5 μm) heat coils with highly tunable resistance based on the thickness of the heating layer (**Fig 3F**, ***supplementary figure 8***). Due to a 1.5 V minimum voltage imposed by the MCU, we chose an optimal resistance of 30 - 32 Ω, requiring 15 - 16 Ω in each of two coils, to maximize heat delivered per unit surface area while preventing PCB brownout. Following polymer channel softening, the grounding spring built into the capsule’s joinable electronic architecture is free to extend, facilitating capsule severance. The complete severing and ejection mechanism is housed within the power capsule (***supplementary table 2***).

Ex-vivo testing of ESCAPE’s severing functionality confirmed its robust ability to sever in the moisture-laden and constrictive GI environment. A polydimethylsiloxane (PDMS) gasket seals the junction between joined capsule segments to prevent moisture ingress for up to 24 hours (***supplementary figure 9***) in acidic conditions. When severed in ex vivo swine intestines mimicking fully fed or empty fasted states, separation times did not increase compared to benchtop controls (**Fig 3G, 3H**).

Beyond facilitating GI transit, ESCAPE’s triggerable severing mechanism can be coupled with a sensing module to enable applications such as closed-loop drug delivery. ESCAPE can load up to 20 mg of powdered drug between its two chambers. Once severed, the payload diffuses into the surrounding fluid and tissue within two hours (***supplementary figure 10***). We demonstrated that ESCAPE’s optoelectronic oxygen sensing module can be used to automatically trigger severing at oxygen concentrations of ≤ 1%, simulating the local luminal oxygen environments of colonic tissue. (**Fig 3I, 3J**). Additionally, the capsule remains capable of manual activation via BLE and is automatically programed to sever after 24 hours to prevent bowel obstruction.

### ESCAPE In-Vivo Evaluation

In vivo in swine, we demonstrated the capability of ESCAPE to perform optoelectronic sensing, sever, deliver an upadacitinib drug payload on command, and safely traverse the GI tract (**Fig 4A, 4B**).

**Figure 4:**
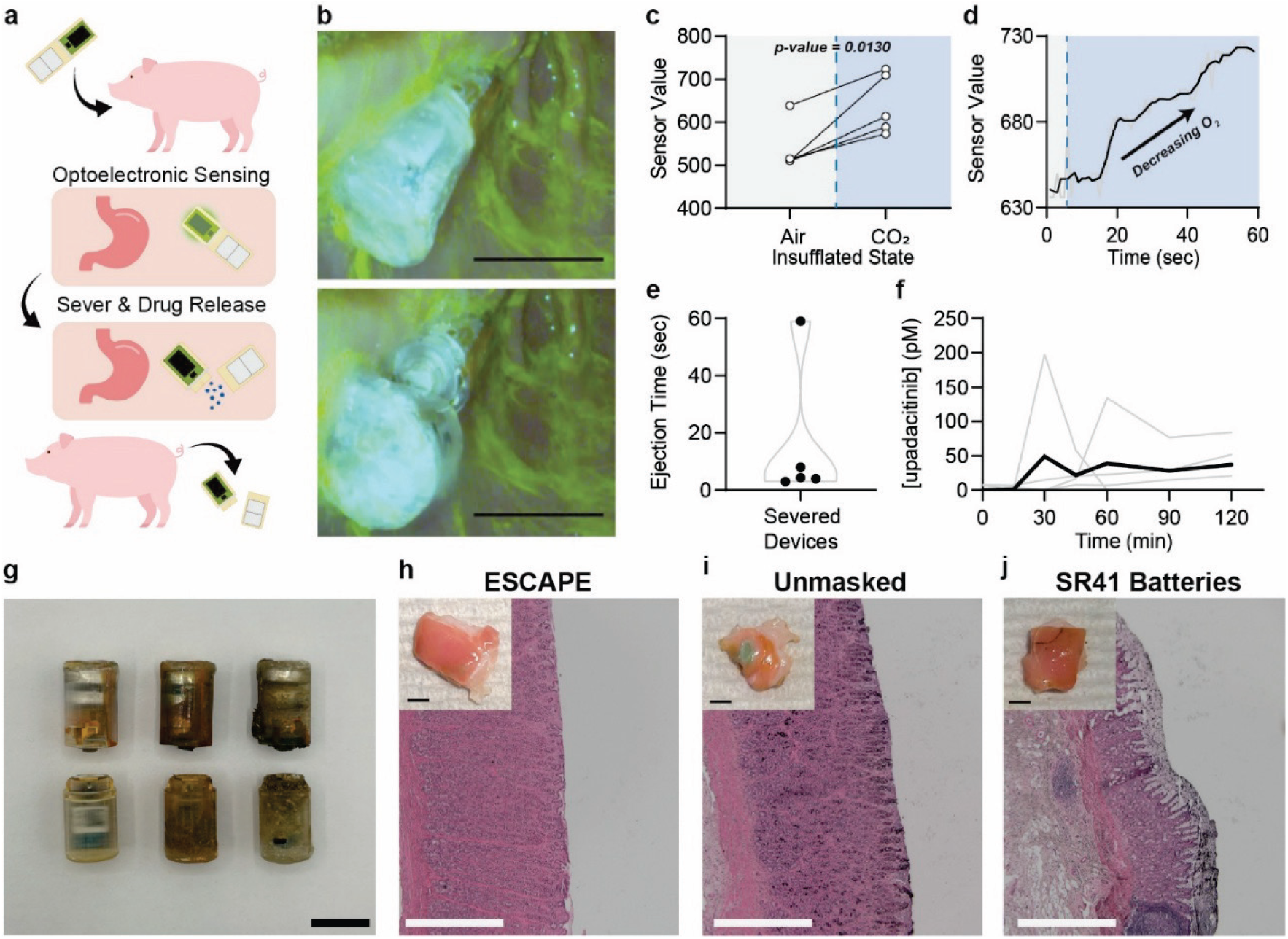
**a,** Experimental flow for in-vivo swine experiments. **b**, Endoscopic images of ESCAPE pre and post severing via BLE triggering in the stomach (scalebar = 10 mm). **c**, Sensor output during air and CO2 insufflation of the stomach (n = 5). Sensor output increases as the oxygen content of the stomach is purged using CO2 insufflation (methods). **d**, Representative real time sensor output as a function of time. Demarcation between air and CO2 insufflation indicated by dashed blue line. **e**, ESCAPE severs within 60 seconds of manual activation via BLE when introduced to the stomach (n = 5 devices). **f**, Once severed, ESCAPE delivers 5 mg of upadacitinib in the stomach. Blood plasma levels are shown 120 minutes following delivery, as quantified via LCMS (bold = Avg, n = 4 devices). **g**, Representative devices before delivery (left), retrieved from the stomach following severing (middle), and retrieved after full passage through the GI tract (right). Following GI transit, no damage was observed in any of the retrieved devices (scalebar = 10 mm). **h**,**i,j**, Photographs and histology following exposure of freshly harvested stomach tissue to ESCAPE’s masked electrical connections (**h**), as well as positive controls of unmasked grounding springs (**i**) and unmasked silver oxide batteries (**j**). Scalebar = 5 mm (top left); 1 mm (bottom right). Illustrations in **a** created with Biorender.com and images licensed from Adobe Stock under Adobe For Enterprise License.

Optoelectronic oxygen gas sensing was shown by observing a significant increase in sensor response under CO2 insufflation following an initial luminal insufflation with air. This manual intervention allowed us to mimic anoxic and oxygenated gastrointestinal states in a controlled manner that ensured near-equivalent changes in oxygen levels across studies (**Fig 4C, *supplementary figure 11***). A real-time increase in sensor response was observed within 60 seconds of CO2 infusion (**Fig 4D**). Data was transmitted from the pill via BLE and recorded on a connected cellphone application. Pills were retrieved following the in vivo procedure to reconfirm their ability to sense oxygen gas levels ex vivo following exposure to the live gastrointestinal environment.

ESCAPE’s severability and potential for drug delivery applications were then demonstrated by triggering capsule separation and upadacitinib release. During the assembly process, devices were loaded with 5 mg of powdered upadacitinib and 15 mg of powdered trehalose (***supplementary figure 12***). In this study, we confirmed on-demand severing via BLE through endoscopic visualization. Devices severed within 60 seconds after triggering (**Fig 4E**). Following triggered device severing, blood plasma samples were collected to confirm upadacitinib uptake (**Fig 4F**). The devices were then allowed to pass through the animals’ GI tracts.

During sever events, the heat produced by ESCAPE to trigger ejection of its capsule segments did not cause thermal damage to GI tissues. A COMSOL model was developed to evaluate the transfer of heat from ESCAPE within a constrictive segment of the GI tract (***supplementary figure 13***). Using this model, we show that the localized hotspots generated by capsule severing do not exceed 2 cumulative minutes at 43°C, while 30 cumulative minutes is required to generate tissue damage^31^. We also performed heating studies on freshly harvested swine stomach and small intestine. We placed the tissue in firm contact with ESCAPE’s power capsule segment and activated the severing circuit for 1 minute, overestimating the time and proximity of the tissue compared to a traditional exposure. Resulting Hematoxylin and Eosin (H&E) histology staining provided evidence that the heat generated by severing did not cause tissue damage when compared to negative and positive controls (***supplementary figure 14***).

Post-severing, ESCAPEs capsule segments were observed to safely pass through the GI tract within 2 weeks without causing discomfort or distress to the animal. This is a typical timeline for swine gastrointestinal transit^32^. We confirmed all capsules severed in-vivo (n = 6) passed safely through the GI tract either through direct retrieval or by radiographs taken at 2 weeks post-operation (**Fig 4G**). In all instances, severed capsule segments were found to remain fully intact following passage and experienced no signs of cracking or mechanical damage.

Following passage of the severed ESCAPE platform, the GI tracts of each animal were observed to exhibit no signs of electrical damage due to battery current leakage or electrolysis. Following euthanasia, the entire GI tracts of each swine were butterflied, and no signs of gross damage were observed (***supplementary figure 15)***. To test whether contacts between the ESCAPE’s electronic leads and gastrointestinal tissue generated tissue damage, we harvested fresh stomach samples and placed tissue in firm contact with either (1) a QTC masked grounding spring (**Fig 4H**), (2) an unmasked grounding spring (**Fig 4I**), or (3) unmasked silver oxide batteries (**Fig 4J**). Subsequent visual observation and H&E histology staining showed that ESCAPE’s QTC masked grounding springs did not generate electrical burns or damage. Unmasked grounding springs and silver oxide batteries, which were used as controls but were not present in ESCAPE, did generate gross damage consistent with previous observations of battery ingestion^33,34^. Random tissue sampling and histology of each animal’s GI organs did not show electrical damage, providing evidence that ESCAPE does not damage the GI tract during passage.

## Discussion

The current potential use cases for ingestible electronic capsules are limited by their inherent risk for generating bowel obstructions. ESCAPE addresses this issue by enabling pills to autonomously sever into pieces with sizes clinically proven to mitigate obstruction risks. Here, we demonstrate ESCAPE’s ability to perform optoelectronic oxygen sensing combined with triggerable, localized drug delivery in vivo in swine. Future studies will be required to assess the safety and efficacy of locally versus systemically delivered upadacitinib in gastrointestinal disorders. Our safety testing for electrical and heating tissue damage as well as obstruction generated no morbidities, but further testing in healthy and diseased patients is needed to confirm the safety of our capsule design. Furthermore, our in vivo testing utilized controlled setups to confirm capsule functionalities. Due to highly variable gastrointestinal transit times, localizing experiments to the stomach uniquely provided consistent gas regulation, capsule severance visualization, and pharmacokinetics monitoring. Expanded testing without endoscopic insufflation in additional gastrointestinal organs will provide supplemental evidence for capsule autonomy. ESCAPE is designed as a platform technology to enable the safe separation of commercially available ingestible electronic components. Further testing with new plug-and-play PCBs possessing different functionalities will help demonstrate the versatility of our capsule design. By reducing the risk of bowel obstructions without reducing capsule volume upon ingestion, ESCAPE allows for complex closed-loop sensing and drug delivery protocols while maintaining a size profile comparable with other nondegradable capsules FDA approved for daily unsupervised dosing.

## Methods

### ESCAPE Form Factor Design & Assembly

Capsule shells and caps were designed using SolidWorks 2025 (Dassault Systèmes) and 3D printed using a Formlabs 3B+ printer. High Temp V2 resin was selected due to its similarity in stiffness and thermal resistance compared to polyether ether ketone (PEEK), a high-performance thermoplastic with known biocompatibility^35^. Both the power and control capsule shells, including all components dedicated to severing, are easily compatible with large scale manufacturing techniques such as injection molding, to facilitate future clinical translation. Polymer interlocks are loaded into the power capsule by applying a melted mixture of polycaprolactone (PCL) and small percentages of polyethylene glycol (PEG) into each polymer channel and allowing it to cool.

Capsule assembly is achieved through a process as shown in ***supplementary figure S12***. First, the empty control capsule’s gasket is lubricated with food grade mineral oil (US+). The control capsule is then inserted into the fully assembled power capsule with its polymer interlocking ridges aligned to the press-fit contact grooves. The control capsule is then twisted 90 degrees to engage the polymer interlock and align each capsule segment. Following this step, the joined form factors’ inner chamber can be loaded with a powdered drug payload by pouring it into the open control capsule. Finally, the PCB can be slotted into the control capsule and sealed with its accompanying cap using medical grade UV epoxy (Loctite 4311).

### Gastric Obstruction Testing

An ex vivo rig was built to evaluate the risk of gastric obstruction in capsules with different dimensions. A peristaltic pump was used to push simulated chyme through tubing wrapped in small intestine tissue (inner diameter 13.8 mm). The peristaltic pump was connected to a 3D printed tube via flexible plastic tubing (McMaster Carr). The 3D printed tube was either straight or curved (180° bend) to mimic the different pathways seen in the SI. The chyme was composed of food designated by the FDA as a high-fat meal^36^. In a blender, two eggs, two strips of bacon, two slices of toast, four ounces of hash browns, and 236.5 mL of water were blended together. The flowrate was set on the pump to provide approximately 4.6 g/min of fluid flow. Obstruction was defined as flowrate being slower than that required to pass 2.5 kg of food within 24 hours (4.167 g/min), which was selected based on an overestimate of average food consumed in one day (2.5 kg/day)^37^. Flow through the empty tube was used as a control. Capsules of varying sizes were secured in the intestinal set up and included 9 × 15 mm, 9 × 26.5 mm, 11 × 15 mm, and 11 × 26 mm sized capsules (diameter × length). The 9 × 15 mm capsule served as a positive control, as it has been designated by the FDA as safe for full GI passage^38^ and the 11 × 26 mm capsule served as a negative control, as it has shown obstruction in 1.4% of patients^18^. We performed extensive characterization of the relationship between obstruction risk and capsule size in healthy bowel conditions, but modeling diseased bowel states may provide better characterization of the heightened risk of obstruction in these cases.

The ex vivo obstruction results were validated for the straight tube set-up using COMSOL Multiphysics version 6.3 (***supplementary figure 1***). Obstruction was classified in the same way as the experimental results, with flowrates below 4.167 g/min being considered as gastric obstruction. The intestinal environment was constructed to model a straight segment of the SI when undergoing a peristaltic contraction^39^. The diameter was set to 15 mm with a tube length of 60 mm. Chyme was sent through the inlet at a rate of 4.6 g/min (incompressible flow, no-slip condition), selected based on our ex-vivo experiments, and was modeled with a density of 970 kg/m^3^ and dynamic viscosity of 0.1 Pa·s^40^. To model the possibility of a GI obstruction, the capsule was introduced with the fluid-structure interaction (FSI) multiphysics interface using fixed capsule boundaries. Capsule diameter and length varied to test capsules of sizes 9 × 15 mm, 9 × 26 mm, 11 × 15 mm, and 11 × 26 mm. Capsule position was changed based on capsule size to ensure all capsules started 25 mm away from the inlet and 0.2 mm away from the intestinal wall. A rectangular outlet was added 28 mm away from the inlet, with a width of 50 mm, depth of 10 mm, and height of 4 mm to mimic backflow and chyme build-up during obstruction. The rectangular outlet was positioned directly opposite the capsule. Both outlets were set to have a static pressure of 0 Pa (intestinal outlet and rectangular outlet). The model was tested without the capsule to ensure the rectangular outlet was not confounding the true outlet flow of the intestine. A flowrate of 4.59 g/min was seen to leave the intestinal outlet, thus indicating no effect of the rectangular outlet in the absence of obstruction. The simulation was run as a time-dependent study, requiring 1 second to reach steady state, and the values recorded were the steady state results.

### ESCAPE Joinable Electric Connections

An fPCB was designed (KiCAD 9.0) and fabricated (JLCPCB) on a two-layer 0.11 mm polyimide substrate to form conformable electric pathways that consume minimal volume within the power capsule segment (***supplementary figure S2***). The fPCB houses press-fit contact rails to connect the control capsule segments rigid PCB to power and the sever circuit. Press-fit contact rails were designed using rectangular electrode pads (dimensions) with 2 U” gold plating and were housed in grooves built directly into the power capsule. Once wrapped around the capsule’s inner diameter, the gently curved geometry of each press-fit rail induces localized stresses when placed in contact with a 90° edge. These localized stresses therefore perform a contact wipe and can create a stable electronic connection even when weakly pressed in contact with the edge of another electrode, which is required for low resistance sliding connections. Sever circuit connections and battery connections were made using through-hole (THT) connections that can be soldered to a curved surface using silver epoxy (MG-8330D). Medical grade UV epoxy (Loctite 4311) is used to fix the fPCB once placed inside the capsule.

The ground connection is formed using a custom spring-loaded electrode pad. A beryllium copper (BeCu) contact pad (Digikey) is directly connected to ground terminal of two SR41W batteries (Seiko) soldered in series using a 32 AWG wire. Silver oxide batteries were chosen due to their enhanced safety profile compared to lithium-ion equivalents. SR41W batteries were specifically selected due to their capacity and high drain designation. They maintain high current pulses required by the optoelectronic sensor over a 24 hour period and tolerate high current loads experienced during sensing and severing functions. A compression spring (Century Spring, 50089SCS; free length = 5 mm, solid height = 1.88mm, OD = 5 mm, k = 1.334 N/mm, 316 stainless steel) provides both pre-load for the grounding connection and mechanical force to eject each capsule once severed. The spring is masked from both the contact pad and the batteries using a 3D printed plunger and housing (Formlabs). All soldered connections were masked with UV epoxy. Battery stacks were masked with both parylene C (PDS 2010, SCS; thickness = 2 µm) and UV epoxy during capsule assembly.

To prevent battery current leakage from the ground connection, a quantum tunneling composite or QTC (Zoflex, ZL45A, 0.5 mm thick) is soldered directly to the ground connections contact pad with silver epoxy. QTC blocks were cut to roughly 1.75 mm long squares using a razor blade before being bonded to the 3D printed plunger with UV epoxy to eliminate small openings for potential tissue contact.

### ESCAPE Joinable Electronics Characterization

Both the press-fit connections and grounding spring were mechanically and electronically assessed using a Mark-10 Tensile Tester in conjunction with a digital multimeter (DMM6500, Keithley). For each press-fit connection, the mechanical resistance of separating the rigid PCB from the press-fit rails was collected (Travel speed: 60 mm/min), in combination with the electrical resistance of the connection during severing. For the ground connection, an ingestible PCB (described in the next section) was pushed into the grounding spring before being quickly pulled off (Travel speed: 60 mm/min). The normal stress of the rigid PCB edge-plated face against the grounding spring required to enable switching between low and high resistance states was assessed by measuring changes in voltage.

### ESCAPE Ingestible PCB

The ingestible PCB was fabricated (PCBWay) on a standard four-layer 0.8 mm FR4 substrate with immersion gold (ENIG) surface finish, measuring 8.1 mm × 12.8 mm × 1.2 mm (***supplementary figure S3***). The DC-DC converter (XCL101A501), an nRF52 MCU (BMD-350) and two 0402 capacitors were soldered on the top side. The optical sensor (MAX86916), two LDOs (S-1317A18 and TPS72748), an NMOS transistor (CSD17381F4) used as a switch in the heating circuit, magnetic sensor (MMC5633NJL), board-to-board (B2B) connector (BM46B-12DP), three 0201 resistors for communication, and three 0201 capacitors were soldered on the bottom side. An overview schematic of ESCAPEs electronics is presented in ***supplementary figure S4***. Three edge-plated pads were used for battery (VBAT), sever circuit (VPCL) and ground (GND) connections. The DC power supply (Keysight E36312A), power profiler kit (Nordic PPK2) and oscilloscope (Analog Discovery 3) were used to validate the PCB functionality and characterize its power consumption in different modes of operation. The nRF Connect application on the mobile device was used to communicate and control the PCB.

### ESCAPE Configurations

The nRF52 MCU on the PCB was programmed using an nRF52 evaluation board and SEGGER Embedded Studio. The PCB was connected to the evaluation board and the firmware with various functionalities was uploaded to the MCU. The firmware enables selection between real-time and closed-loop operating modes, independent configuration of optical gas sensing parameters such as LED current (1 mA - 50 mA), sensing interval (1 - 10 s) and O₂ concentration threshold value for closed-loop operation, and control over enabling or disabling the heating circuit. In real-time mode, the device performs gas sensing using predefined LED current and sensing interval parameters, transmitting the data to a mobile device via BLE, where it is displayed in real time on the nRF Connect app. Upon reaching a target O₂ level, the heating circuit can be manually activated to trigger separation. In closed-loop mode, the device also continuously monitors O₂ levels using predefined parameters for LED current and sensing interval. Instead of waiting for a manual command from the mobile device, the MCU automatically activates the heating circuit once a user-defined threshold, set at the beginning, is exceeded. Aside from these modes, the system also supports an automated sensing mode. Once the device is powered by the batteries, the user can enable this mode, where sensing is performed using user-defined LED current and sensing intervals. However, instead of transmitting data over BLE, all measurements are stored locally on the MCU. This functionality is particularly useful in cases where BLE connectivity is unreliable. After the procedure is completed and the pill is retrieved, the user can reconnect to the device over BLE and download all data stored locally on the MCU. A total of 3600 sensing data points can be stored on the MCU, corresponding to 1 hour of operation at a 1 s sampling interval, 10 hours at 10 s, and 60 hours at a 60 s sampling interval. Finally, the firmware runs a low-power timer that triggers the heating circuit 24 hours after the device is powered on, ensuring device disassembly regardless of sensing values and safeguarding against potential measurement inaccuracies.

Irrespective of firmware selection, the ingestible PCB performs sleep cycling of its electronics to minimize power consumption during operation (***supplementary figure 5***). Once the PCB is powered, the MCU starts BLE advertising to indicate that it is ready to connect to a mobile device. During this period, the optical sensing module and the low-dropout regulator that supplies the LEDs are kept in sleep mode. Since BLE advertising itself is relatively power-hungry, the firmware alternates a 5-second advertisement window with a 5-second sleep cycle, during which only a single low-power timer remains active. Upon activation of optoelectronic sensing, the PCB likewise briefly activates both the optical module and the low-dropout regulator at user-defined intervals before immediately returning both to sleep mode until the next sensing cycle.

### Optical Gas Sensing Membrane Fabrication

A combination of Platinum octaethylporphyrin (PtOEP, Sigma Aldrich) and Coumarin 545T (C545T, Fisher Scientific) was used as the luminophore indicator for the oxygen gas sensing membrane. To prepare a stock solution, 30.3 mg of PtOEP and 52.7 mg of c545T were added to 10 mL of toluene in a glass bottle and sonicated until fully dissolved. In a separate glass bottle, 10 g of Sylgard 184 polymeric base (Dow Corning) was combined with 2 mL of the stock solution. This mixture was vortex mixed until a uniform consistency was achieved. Subsequently, 1 g of Sylgard 184 curing agent was incorporated, and the mixture was vortex mixed again. For membrane fabrication, the resulting solution was drop-cast onto a clean, flat, 80 mm diameter glass petri dish to achieve the desired thickness. The solution was degassed under vacuum to remove air bubbles. The membrane was then cured at 60°C for 4 hours.

### ESCAPE Optoelectronic Sensing Characterization

To accurately relate sensor output to oxygen concentrations, a calibration curve must be generated by recording sensor output at known oxygen concentrations. We then compared the ability of the optoelectronic sensing module to function when enveloped by ex-vivo swine small intestine generating similar calibration curves for comparison with ambient calibrations. Finally, optoelectronic components that were tested in-vivo were stored and recalibrated to evaluate the sensor module’s ability to survive an in-vivo gastric environment.

All calibrations were conducted in a vacuum chamber connected to a CO₂ line and a vent line. Devices were either placed in the chamber under ambient conditions or inserted into an empty ex-vivo section of swine small intestine (Kinship Butchers) and partially submerged in PBS (Thermofisher). The O₂ concentration inside the chamber was manipulated by sequential infusions of CO₂ followed by venting to reduce pressure and remove excess air, which naturally is displaced to the top of the chamber by heavier CO2 gas. A commercial O₂ detector (Forensic Detectors, FD-90A-O2) was used as a reference sensor in all experiments. All sensing trials were conducted in complete darkness.

For integration with optoelectronic sensing, the standard control capsule cap was modified to incorporate a gas-permeable membrane (SSPM823-005-12″) sealed to its inner surface with UV epoxy. Gas permeable membranes were installed at the base of each control capsule segment cap. This location was chosen to minimize tissue contact in tightly constricted scenarios, where the joined capsule naturally aligns with the tubular geometry of the GI tract. When aligned with the intestinal lumen, tissue conformally covers any slots built into the capsule’s cylindrical face, which we found to significantly inhibit gas from infiltrating the capsule’s inner chamber. (***supplementary figure S6***).

Closed-loop trials were conducted using the same experimental set-up as standard sensing trials. Once inserted in the vacuum chamber, the oxygen concentration was lowered from an ambient level to 1% to collect the corresponding threshold value for that sensor. The vacuum chamber was then quickly brought back up to ambient oxygen concentrations and then continuously pumped down at a slow, constant rate to assess closed-loop actuation. Three devices fired prematurely due to this time, for reasons unrelated to the test, and they were not included in the final dataset. One device fired because the light was turned on during the middle of the test, causing ambient interference. Two devices experienced unexpected spikes in sensor value immediately as the CO2 was turned on due to a rapid influx of gas, causing premature actuation to occur. These devices were excluded from the sample set.

### Resistive Heater Design & Assembly

Flexible thin film heaters were fabricated by evaporating 300 nm of Aluminum (Evaporation rate: 3 Å/sec) followed by 10 nm of Gold (Evaporation rate: 0.5 Å/sec) onto 1 mil polyimide sheets (Digikey) under strong vacuum. A femtosecond laser-cutter was then used to pattern (4W, 60 kHz, 10% power, no repetitions, OPTEC WS-Flex USP) and cut (4W, 60 kHz, 30-35% power, 2 repetitions, Optec) each resistive heating coils (***supplementary figure 8***). Each coil was then secured to the curved inner diameter of each polymer channel using UV epoxy and soldered to the fPCB with silver epoxy (MG-8330D) to form the full sever circuit (two coils soldered in series along one pathway). All soldered connections were masked with Loctite 4311.

The electrical resistance of each coil can be determined analytically using the equation:

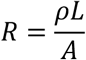

Where ρ is the resistivity of the material, L is the length of the coil, and A is its cross-sectional area. The thickness of the coil disproportionally impacts its resistance through both its impact on cross-sectional area A and resistivity ρ. Thin film evaporated metals are known to exhibit increased resistivity due to the formation of grain boundaries and electron scattering^41^ Compared to a bulk resistivity of 2.65 to 2.82 × 10^−8^ Ω·m, our fabricated coils exhibited measured resistivities of 4.80 × 10^−7^ Ω·m when tested. Consequently, coil length and spacing were fixed at 200 µm and 50 µm to maximize the surface area of the heating region, and the thickness was varied to produce coils of varying resistance (**Fig 3F**).

Aluminum was chosen as a material choice due to its common availability and relative biocompatibility. A thin gold layer (10 nm) was used to mask the main heating layer to prevent long term resistance changes due to oxidation.

An optimal resistance of 15 - 16 Ω was determined for each individual heat coil to maximize heating efficiency under the power constraints of ESCAPEs electronic components. The MCU on the PCB requires a constant supply of 1.55 – 1.6 V in order to operate, including when current is flowing through the heating circuit. During a severing event, the SR41W batteries are allowed to freely discharge across the heating circuit, reducing the operating battery voltage proportional to the applied resistance as defined by Ohms Law, Voltage = Current * Resistance, where the batteries are allowed to approach their peak discharge current. Additionally, the batteries experience rapid voltage sag when discharging due to depleting battery capacity. The resistive heater must therefore have sufficient resistance to maintain 1.55V when the sever circuit activates and account for voltage sag over the duration of the severing window. As a result, we chose a total target resistance of 30 - 32 Ω to maximize the power we deliver across the resistive heating regions while ensuring that MCU failure will not occur over a sever window > 1 min. Each individual coil provides 15 - 16 Ω across its active heating surface before being connected in series to achieve the final threshold with minimal resistance increase during the soldering process.

### ESCAPE Watertight Characterization

To make the junction between each capsule segment watertight, a gasket was fabricated from 10:1 PDMS (Sylgard 184). Following mixing, the PDMS was spin-coated on a glass slide to produce thicknesses of 350 ± 50 um (300 rpm, 30 seconds, 300 rpm ramp). PDMS was allowed to cure at 65C for 1 hour under vacuum. The gasket was then laser-cut using a femtosecond laser (4 W, 60 kHz, 70% power, 60 repetitions, Optec). UV epoxy was finally used to bond the gasket on one side to the control capsule segment (***supplementary figure S9***)

To test the devices’ ability to prevent moisture infiltration prior to severing, we measured the change in mass of capsules after submerging in simulated gastric fluid (SGF, described in a later section) for 24 hours. For watertight testing, dead SR41W batteries and 3D printed mock PCBs with similar dimensions and weight were loaded in lieu of active electronics. Over 24 hours, the devices were monitored in simulated gastric conditions (37°C, 500 rpm spin rate) for visual signs of leakage, as we loaded the capsules with methylene blue that could be detected in the bulk fluid if it leaked. After 24 hours, devices were removed, patted dry, and weighed to evaluate the extent of moisture penetration via net mass change.

### ESCAPE Severing Characterization

The mechanical pull force that the loaded dovetail channels can resist was first tested using a Mark10 Tensile Tester. The ultimate tensile strength of each dovetail channel was first assessed by tensioning each capsule seal with the tensile tester (Travel speed: 1 mm/min) until failure occurred. Once the ultimate tensile strength of the dovetail channels was determined, channels were tested under cyclic loading conditions (Travel speed: 60 mm/min) at peak tensile forces ranging from 4.5 – 5 N to evaluate the seals durability. Devices used in mechanical testing were assembled with heat coils wired to a power source without all other accompanying electronics.

As shown in ***supplementary figure 7***, a dovetail geometry was selected for the polymer channel interlocks due to significantly increased ultimate tensile strength and decreased yield strength variability compared to rectangular channels. This is due to the slanted dovetail design’s engagement of the casted polymer’s tensile modulus, whereas a rectangular channel chiefly holds back force using the shear modulus of interface between the surface of the channel. In particular, a dovetail channel measuring 1 mm in length provides equivalent ultimate tensile strength to a 4 mm long rectangular channel with the same width as the short base of the dovetail. Dovetail geometries are also non-complex and do not present additional manufacturing burden compared to straight channels. We therefore chose a 1 mm long dovetail design for ESCAPE to minimize the polymer volume cast into the channels and its resulting thermal mass.

Full capsules were then assembled and severed in room temperature, benchtop conditions. In benchtop trials, SR41W silver oxide batteries were first drained for 24 hours at a constant current draw of 0.61 - 0.615 mA of current to simulate worst case power demands from capsule electronics. The power dissipated by the heat coils during severing was measured using a digital multimeter.

Ex-vivo severing trials were conducted in both an empty swine SI and SI filled with blended FDA high fat diet to simulate a fed state. Tested devices were assembled and inserted into either an empty or fed SI model and partially submerged in PBS at 37°C prior to severing. To simulate the depleted battery performance measured in the benchtop trials, a programmable power supply (Siglent SPD3303X) was used to mimic the power dissipated over a 1-minute interval. Severing was confirmed with visual indication of a crease created in the SI and by post-trial removal of each separated capsule segment.

### ESCAPE Drug Expulsion Testing

To assess the potential for capsule severing as a triggerable drug release mechanism, capsules were assembled as previously described for watertight characterization testing with the addition of 5 mg of methylene blue loaded into the capsule’s free chamber space (***supplementary figure S10***). Assembled capsules were submerged in 50 mL of water, which was selected to match the maximum detectable concentration of methylene blue (0.1 g/mL) with UV-Vis, at 37°C. Capsules were left submerged for 10 minutes before severing to collect a control measurement and confirm that the assembled device was not leaking drug prior to severing.

Following severing via heat-gun (Seekone), both capsule segments were left submerged for 120 minutes. During that time, two 150 µL aliquots were sampled at t = 0 sec, 5 sec, 15 sec, 30 sec, 45 sec, 60 sec, 5 min, 20 min, 60 min, and 120 min were sampled. Aliquots were later analyzed using a plate reader (BioTek Synergy H4). Collected samples were compared to a standard curve with dilutions up to 0.10 g/mL of Methylene Blue to measure the %dose that diffused out of the capsule during each sampling interval.

### In Vivo Testing

All procedures were reviewed and approved by Emory University’s Institutional Animal Care and Use Committee (STUDY IPROTO202400000099). All in vivo studies were performed at Emory University on female Yorkshire swine weighing 30 – 40 kg (Oak Hill Genetics). Animals were placed on a liquid diet for 24 h before the study and were fasted overnight before the study. We anesthetized the swine with intramuscular injections of TKX (telazol 4.4 mg/kg, ketamine 2.2 mg/kg, and xylazine 2.2 mg/kg) and intubated and maintained on 1 – 3% isoflurane in oxygen. Devices were then delivered orally to the stomach via orogastric tube and imaged using PENTAX EPK i7010 to visualize the stomach with the animal in the left lateral position.

The optoelectronic sensing module was assessed in the gastric environment through the use of air and CO2 insufflation to either infuse or purge the stomach with oxygen. Prior to oral delivery, a mobile device was connected to the capsule and used to initiate its optoelectronic oxygen sensing module’s automated sensing mode. BLE was then disconnected prior to device delivery. Next, the activated device was introduced to the gastric environment and insufflated with air via the endoscope. Once baseline measurements were collected, the stomach was then partially deflated via endoscope and quickly re-insufflated with CO2 insufflation bags connected directly to the endoscope (PENTAX EPK-i7010). Each bag was filled with 750 mL of CO2, which was introduced to purge oxygen gas from the stomach. 1 - 2 bags were administered to the animal. Following a 5 - 10-minute measurement interval, devices were retrieved from the animal and reconnected to BLE to transmit sensing data directly to a mobile device. The stomach of the animal was also deflated to prevent unwanted gas from propagating through the lower GI tract.

We observed changes in sensor response between ambient measurements taken before and after introduction to the gastric environment (***supplementary figure 11***). The observed drift in sensor response does not appear to impact the device’s response to fluctuations in oxygen concentration and may be due to biofouling and subsequent light scattering during sensing.

To demonstrate ESCAPE’s severability, capsules were assembled with a pre-mixed combination of 5 mg upadacitinib (Tocris Bioscience) and 15 mg of Trehalose (Fisher Scientific) to serve as a filler excipient. Once capsules were assembled, BLE connection with a mobile device was established and the device was delivered to the stomach via an overtube. Prior to severing, an initial blood plasma sample was collected for a t = 0 timepoint. Severing was then initiated manually and visualized via the endoscope. All devices severed at similar timescales within one minute, similar to ex vivo testing (**Fig 4E**). One observed device exhibited a 1-minute delay in severing due to a poor BLE connection that needed to be reestablished. Following severing, plasma samples were collected at 15, 30, 45, 60, 90, and 120 min to correspond with previously reported peak plasma concentrations of upadacitinib formulations^42^. Samples were then analyzed using Liquid chromatography–mass spectrometry (LCMS) with Pexidartinib (Fisher Scientific) being used as an internal standard^43^. During LCMS measurements, samples were 10x concentrated to ensure detection within single nM concentration ranges.

### Validation of ESCAPEs ability to facilitate GI transit

Following delivery and severing of ESCAPE capsules, capsule segments were allowed to pass through the GI tract over a period of two weeks. Daily cage examinations were conducted to observe animal stool for passed devices and animal behavior for signs of discomfort. In addition, radiographs were performed at 1 and 2 weeks post severing to visualize capsule passage through the GI tract. A total of n = 6 delivered devices were confirmed to safely pass out of animals through a combination of device retrieval in stool and post-operative radiographs to confirm the absence of devices in the GI tract. All devices that were dosed to the animal were retrieved intact.

14 days after the conclusion of in-vivo experiments, animals were euthanized and their GI tracts were harvested and butterflied as shown in ***supplementary figure 15***. During the butterflying process, the GI tract was examined and photographed to search for gross damage caused by capsule segment passage. No damage was found. Representative tissue samples of the stomach, small intestine, and colon were harvested for Hematoxylin & Eosin (H&E) histology staining to provide further evidence that ESCAPE does not distress the GI tract during passage. Again, no tissue damage was found. After one of our in-vivo studies, but before the pill passed, one animal was euthanized due to health complications unrelated to any endoscopic experimental procedure. Following euthanasia, intact capsule segments were retrieved from the stomach and small intestine. The GI tract of the animal was grossly assessed for damage, and no damage was found as determined by a veterinarian and a pathologist. It was not possible to completely butterfly the complete GI tract of this swine due to the timing of its euthanasia.

### Thermal and Electrical Safety Characterization

The transfer of heat generated by ESCAPEs severing event was first modeled using COMSOL Multiphysics version 6.2. To maximize the validity of the model, 3D models of the capsule and heating coils were simulated to deliver the power curve developed in benchtop severing experiments (see previously) over the course of 60 seconds to mimic a worst-case heat exposure scenario (***supplementary figure 13***). Built-in material models with thermal conductivities were used for the following materials: PEEK (κ = 0.366 W/(m·K), Cp = 1340 J/(kg·K)), polycaprolactone (κ = 0.2 W/(m·K), Cp = 2000 J/(kg·K)), and polyimide (κ = 0.150 W/(m·K), Cp = 1100 J/(kg·K)). A simple approximation of the intestinal environment was modeled with a 19 mm OD × 5 mm thick × 15 mm long body of water that surrounded the outer diameter of the power capsule segment. A thickness of 5 mm was defined by referencing literature for narrow ID sections of the SI in humans^44^. A flowrate of 1.16 × 10^−7^ m^3^/s was defined for the surrounding aqueous environment to simulate chyme flow in the SI using the laminar flow physics model (incompressible flow, no-slip condition) and the initial temperature at t = 0 was set to 37°C. To model a sever event, a boundary heat source of 37602.77145 W/m^2^ was applied to the model of the heat coil surface in the capsule power segment for 60 seconds using the Heat Transfer in Solids and Fluids module. This value corresponds to the average power dissipated during a minute long heating cycle collected during benchtop sever testing. The simulation was run for a total of 30 minutes following a one minute sever event to study heat dissipation over time in simulated physiological condition. During and following the one-minute severing event, a 3D cut point was defined at the centered intersection between the capsule segments outer diameter and the aqueous environment and recorded for all simulation time-points.

Following analysis via the computational model, ex vivo heating experiments were performed by activating the heating circuit in an ESCAPE power capsule segment when pressed against either ex-vivo swine stomach or small intestine (***supplementary figure S14***). Treated tissues were first placed in a PBS bath that was maintained at 37°C via a hotplate. Heating was then performed as previously described for ex-vivo severing testing. Post-heating, tissues were then analyzed using H&E histology staining and compared to both negative controls left at 37°C, and positive controls that were heated for 10 minutes at 60°C through direct contact with a hotplate heating surface. Histology tissue samples were fixed overnight in 10% formalin buffer (VWR), then dehydrated by an automatic tissue dehydration system. The dehydrated tissue then was embedded in paraffin and then 5 μm thick sections were obtained with a rotary microtome (Leica Biosystems). H&E staining was then performed with an automatic stainer (Leica Biosystems). Stained slides were imaged using a slide scanner (BioTek Cytation 7; Agilent). Comparison between the tissue treated with ESCAPE versus negative controls and positive controls did not yield any visual indications of tissue damage.

During visualization of the GI tract following swine euthanasia, freshly harvested stomach tissues were assessed for electrical damage (***supplementary figure 15***). Additionally, we artificially created preferential opportunities for electrical damage by placing tissue in direct contact with either (1) QTC masked power capsules (what was used in ESCAPE), (2) unmasked power capsules (not used in ESCAPE), or (3) 2 unmasked SR41W batteries to serve as a positive control for electrical tissue damage (**Fig 4H - 4J**). No powering unit groups were drained prior to being placed in contact with ex-vivo stomach tissue. After thirty minutes of exposure, treated tissues were photographed and later analyzed with H&E staining. The photographs of the unmasked treatment groups were compared to butterflied GI organs to search for sites of electrical damage and histology samples were likewise compared to representative histology of each GI organ. No damage was found on any representative histology sample.

### SGF Preparation

Simulated gastric fluid was prepared as described by Pan et al. 3 g of NaCl was dissolved in 1450 mL of deionized water^45^.The pH was adjusted with diluted HCl to 1.2 ± 0.1. Deionized water was used to adjust the acidic solution to a final volume of 1500 mL.

## Data analysis

Recorded data was parsed using custom scripts written in Python using numpy, pandas, and matplotlib libraries and in MATLAB R2025A (Mathworks). Graphical representations of data were generated using GraphPad Prism 8.4.2. Statistical analysis was performed in GraphPad Prism 8.4.2 and R 4.3.5 using the readxl, tidyr, and multcomp libraries.

## Acknowledgements

We thank all members of the Abramson Lab for their discussions and insights. We thank Allison Hayward, Crystal Gergye, Andrey Krasnopeyev, Alex Bailey, and Robin Michelle Carter for their help with animal experiment design and training. We thank the Parker H. Petit Institute for Bioengineering and Bioscience, School of Chemistry and Biochemistry, and the Office of the Executive Vice President for Research for their generous support. This work was in part conducted at the Georgia Tech Institute for Matter and Systems, a member of the National Nanotechnology Coordinated Infrastructure (NNCI), which is supported by the National Science Foundation (Grant ECCS-2025462).

## Author Contributions

S.H. and A.Abr. conceptualized and designed the experiments supporting the development of ESCAPE. A.Abd. designed and fabricated the ingestible PCB for ESCAPE. S.H. designed ESCAPEs joinable and severable design. S.H., J.Z.D., R.G., X.M., S.P., M.C.L. fabricated ESCAPE components. A.Abd. and M.S.I. fabricated, tested and characterized the optoelectronic gas sensing membrane. S.H., A.Abd., M.C., J.Y.C., X.M., S.P., M.C.L., and J.P. performed benchtop and ex-vivo experiments for the characterization of ESCAPEs electronic and severable functions. S.H., Z.L., J. Z. D., J, J., and A.Abr. performed or assisted with the in vivo experiments endoscopically delivering ESCAPE in swine. A.Abr. and Y.K. provided funding and supervised the project. All authors contributed to writing and editing the manuscript.

Conceptualization: SH, AAbr

Methodology: SH, AAbr

Investigation: SH, AAbd, MC, JYC, MSI, ZL, JZD, JJ, RG, XM, SP, MCL, JP

Visualization: SH, AAbr, AAbd, YK

Funding Acquisition: AAbr, YK

Project Administration: SH, AAbd, AA, YK

Supervision: AAbr, YK

Writing – original draft: SH, AAbd, MC, JYC, MSI, ZL, JZD, JJ, RG, XM, SP, MCL, JP, AAbr, YK

## Diversity, equity, ethics, and inclusion

We support inclusive, diverse, and equitable conduct of research.

## Data and materials availability

All data are available in the main text or the supplementary materials. Materials used in this manuscript are available for purchase from the vendors listed in the methods and supplementary materials. Device fabrication, characterization, and materials synthesis methods are described in the methods section. Devices are available from the authors upon request. An MTA may need to be negotiated with Georgia Tech prior to the use of materials outlined in this manuscript.

## Code availability

The code developed for this study mainly consists of device firmware used for hardware operations. Limited Matlab® and Python scripts were generated for data organization. All code is available upon request.

## Supplementary Materials

Tables S1-S2 Figs. S1-S15

