## Supplementary for "Self-severing circuits facilitate passage of ingestible electronic sensor-guided therapeutics"

### This file contains:

**Table S1: List of PCB components and manufacturers**

| Item # | Designator | Manufacturer | Part # | Description |
|---|---|---|---|---|
| 1 | U1 | Torex Semiconductor Ltd | XCL101A501ER-G | DC DC CONVERTER 5V |
| 2 | U2 | Texas Instruments | TPS72748YFFT | IC REG LINEAR 4.8V 250MA 4DSBGA |
| 3 | U3 | ABLIC Inc. | S-1317A18-A4T2U4 | IC REG LINEAR 1.8V 100MA HSNT4-B |
| 4 | U4 | Memsic Inc. | MMC5633NJL | 30 GAUSS, MONOLITHIC, HIGH PERFO |
| 5 | U5 | Analog Devices Inc./Maxim Integrated | MAX86916EFD+ | SENSOR OPTICAL I2C OUTPUT |
| 6 | U6 | u-blox | BMD-350-A-R | RF TXRX MOD BT TRACE ANT NORDIC |
| 7 | U7 | Texas Instruments | CSD17381F4 | MOSFET N-CH 30V 3.1A 3PICOSTAR |
| 8 | U8 | Hirose Electric Co Ltd | BM46B-12DP-0.35V(53) | CONN HDR 0.35MM SMD GOLD 12POS |
| 9 | C1, C2 | KYOCERA AVX | KGM05CR50J226MH | CAP CER 22UF 6.3V X5R 0402 |
| 10 | C3, C4 | Murata Electronics | GRM035R60J475ME15D | CAP CER 4.7UF 6.3V X5R 0201 |
| 11 | C5 | KYOCERA AVX | KGM03BR50J105KH | CAP CER 1UF 6.3V X5R 0201 |
| 12 | R2, R3 | YAGEO | RC0201JR-072K7L | RES 2.7K OHM 5% 1/20W 0201 |
| 13 | R4 | YAGEO | RC0201DR-071ML | RES 1M OHM 0.5% 1/20W 0201 |

**Table S2: Comparative analysis of ESCAPE with related ingestible technologies**

| Criteria | ESCAPE | Kong et al[1] | Zhang et al[2] | Huang et al[3] | Raman et al[4] | Liu et al[5] |
|---|---|---|---|---|---|---|
| Device Description | Severable Pill | Gastro-retentive Pill | Gastro-retentive Device | Gastro-retentive Pill | Deflatable Gastro-retentive Balloon | Bioresorbable, Shape Adaptive Film |
| Allows Electronics (Yes/No) | Yes | Yes | No | Yes | Yes | No |
| Biocompatibility | High | Medium (10 wt% CNT loading) | High | High | High | High |
| Reported volume during passage | 9 x 15 mm | 9 x 27 mm (estimate) | Partial to complete dissolution | 26.99 x 8.42 mm | 50 to 100% volume reduction of 71 mL inflated balloon | Completely Dissolved |
| Actuation Mechanism | Joule heating of polymer interlock | Joule heating of electroactive adhesive or passive hydrolysis | Passive Enteric dissolution | Induced Enteric Dissolution | Light Degradable Hydrogel | Bioresorbable Material Design |
| Triggerability | Manual triggering via mobile device or pre-programmed conditional triggering | Manual Triggering via mobile device or Passive Hydrolysis | Passive Disintegration; Not Triggerable | Ingestion of alkaline solution | Endoscopic Tethered or untethered light triggering (365 nm) | Passive Disintegration; Not Triggerable |
| Time-Delay | 0-1 min | 1 minute during triggering; up to 30 days for passive dissolution | 0 to 14+ days | 0 to 3 hours, not demonstrated in-vivo | 0 to 6 hours | 0 to 30 days |
| Power Demands for Actuation | 1.43 mWh | Not mentioned | n/a | n/a | 5.19 mW $cm^2$ for 70 min or 11.4 mW $cm^2$ for 30 min | n/a |
| Power Source | 2 silver oxide coin cell batteries (7.9 x 7.2 mm) | Coin cell batteries (not described) | n/a | Rechargable lithium coin cell battery (5.8 x 1.8 mm) | Three zinc-air coin cell batteries (5.8 x 10.8 mm) | n/a |
| Communication Capability | BLE | BLE | n/a | n/a | n/a | n/a |

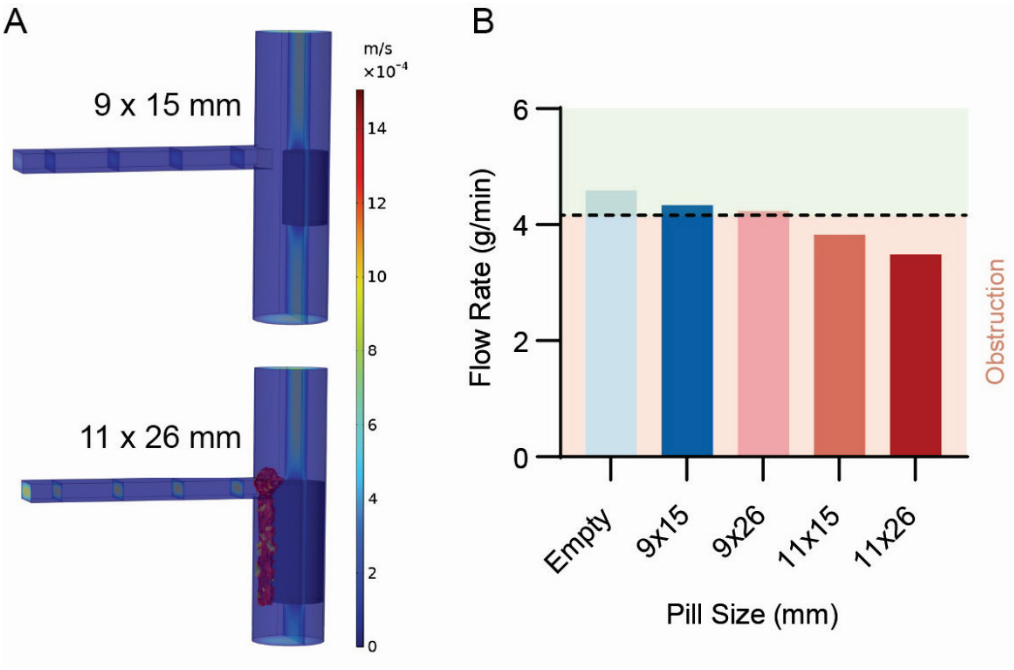

**Fig. S1: COMSOL simulated capsule size-dependent obstruction.** (A) Capsule sizes of 9 x 15 mm (FDA approved size), 11 x 15 mm, 11 x 26 mm (obstruction in 1.4% of patients), and 9 x 26 mm, were tested in a modeled straight segment of the small intestine undergoing a peristaltic contraction (15 mm ID) to observe obstruction. (B) Comparison of flowrates for each capsule size. A dashed line indicates a flowrate below 4.1 g/min is defined as GI obstruction

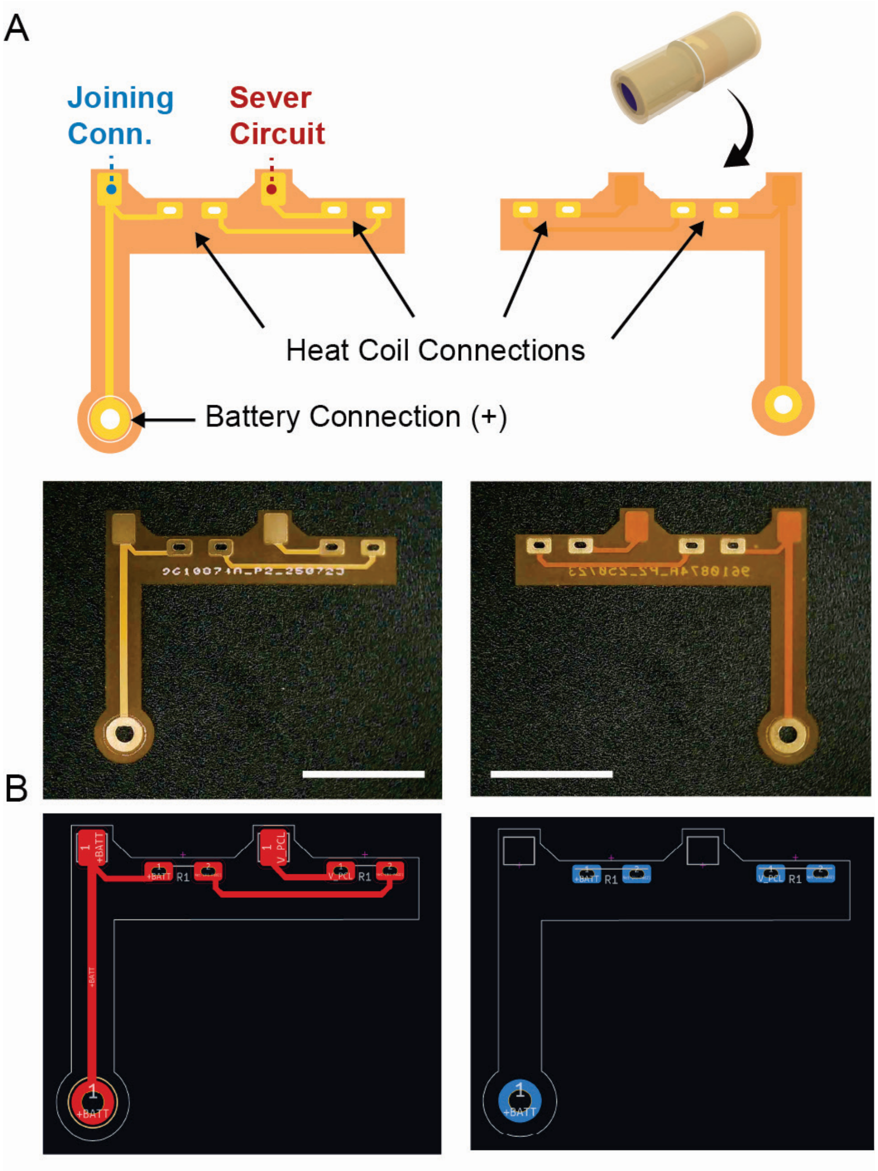

**Fig. S2: Joinable electronics utilize conformal flexible printed circuit boards.** (A) Flexible PCBs containing press-fit contacts and through-hole (THT) contact pads that comfortably wrap around ESCAPE's inner diameter (scalebar = 10 mm). (B) The fPCB's internal wiring connects soldered heating coils and positive battery connections to provide power to the ingestible PCB and house the sever circuits wiring. Left: PCB front. Right: PCB back.

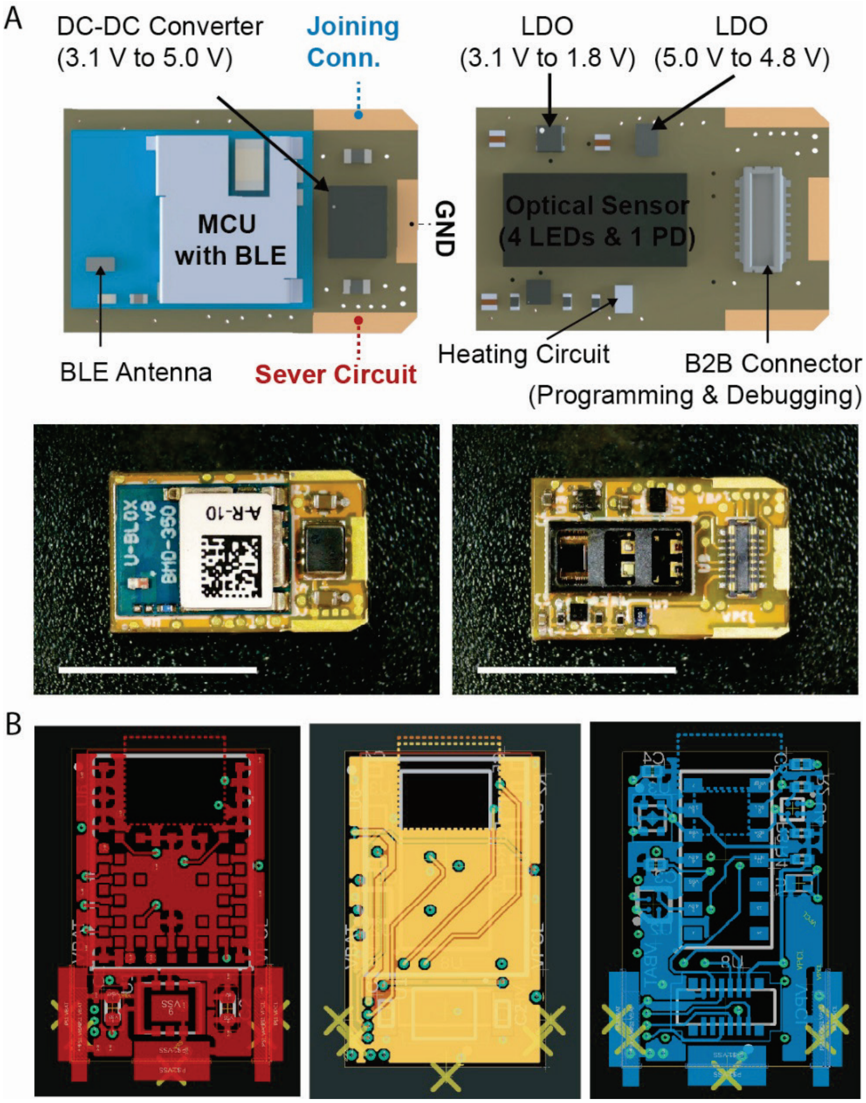

**Fig. S3: Commercial PCBs are compatible with ESCAPE through minimal adjustment to board architecture.** (A) Ingestible PCB implementing three edge-plated contacts without sacrificing space for key components, including: an nRF52 microcontroller unit (MCU), two Low-dropout regulators (LDOs), an n-channel Metal-Oxide-Semiconductor (NMOS) transistor designated as the heating circuit, a board-to-board (B2B) connector, and an optical sensing module (scalebar = 10 mm). (B) PCB fabrication layout illustrates top (left) and bottom (right) routing and power planes as well as two middle ground and minor routing planes (middle).

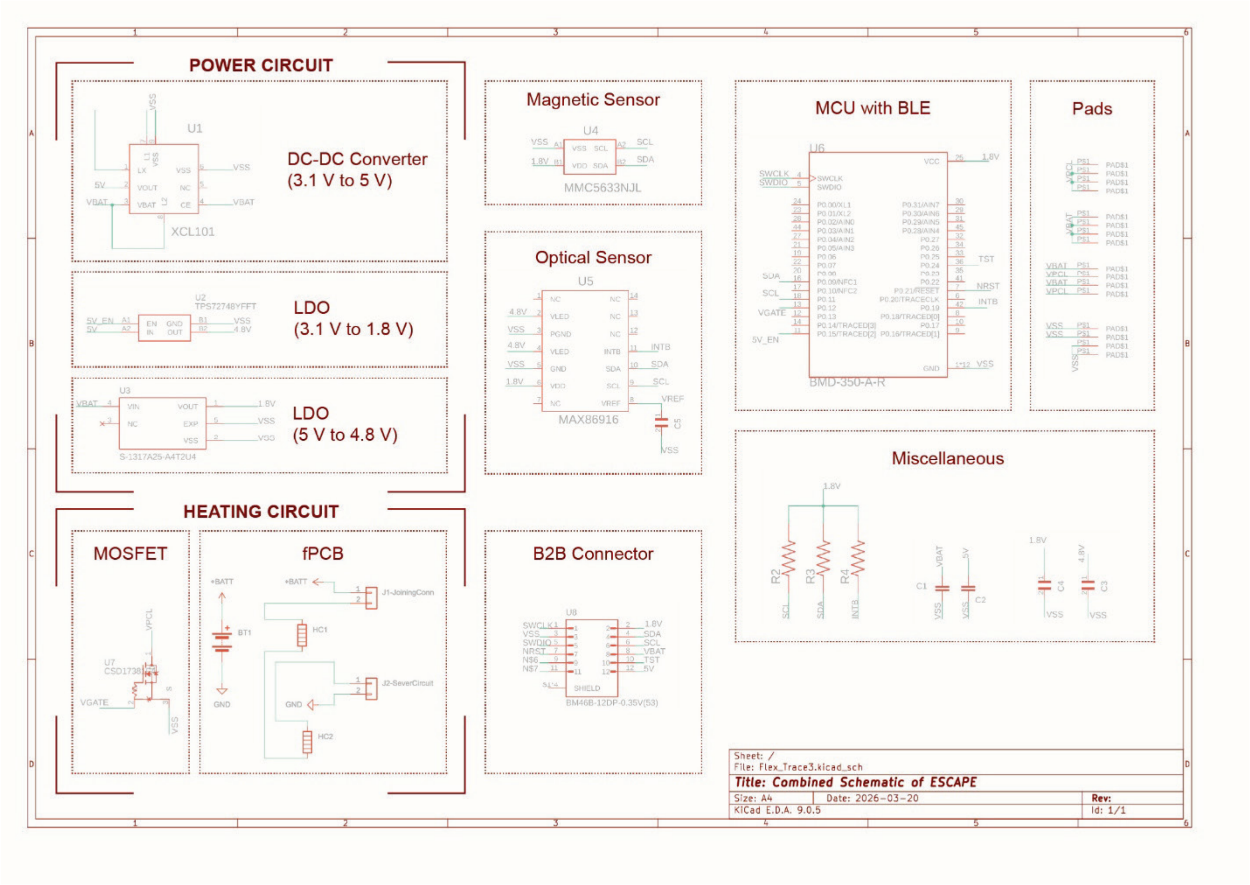

**Fig. S4: Schematic of ESCAPE electronics,** consisting of a microcontroller, power circuit, heating circuit, and different electronic modules. Supporting peripherals and miscellaneous circuitry are also shown.

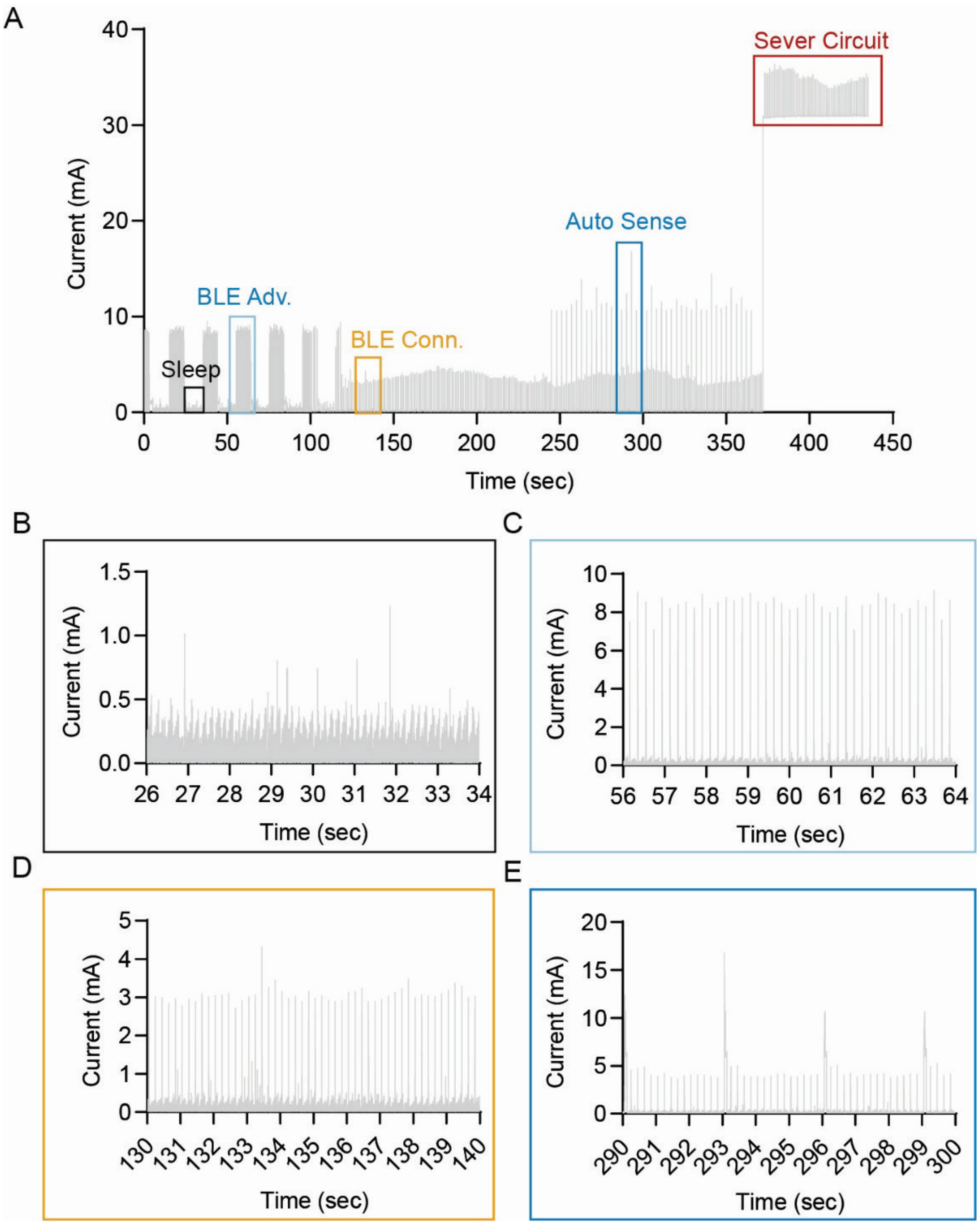

**Fig. S5: Full current profiling of ESCAPE PCB configurations.** (A) Full current profile with highlighted sections corresponding to (B) Sleep ($I_{avg}$ = 85 μA), (C) BLE advertisement ($I_{avg}$ = 201 μA), (D) BLE connection with no functions enables ($I_{avg}$ = 91 μA), and (E) BLE activated with auto sensing enabled ($I_{avg}$ = 253 μA). The sever circuit is activated at the far right ($I_{avg}$ = 31 mA)

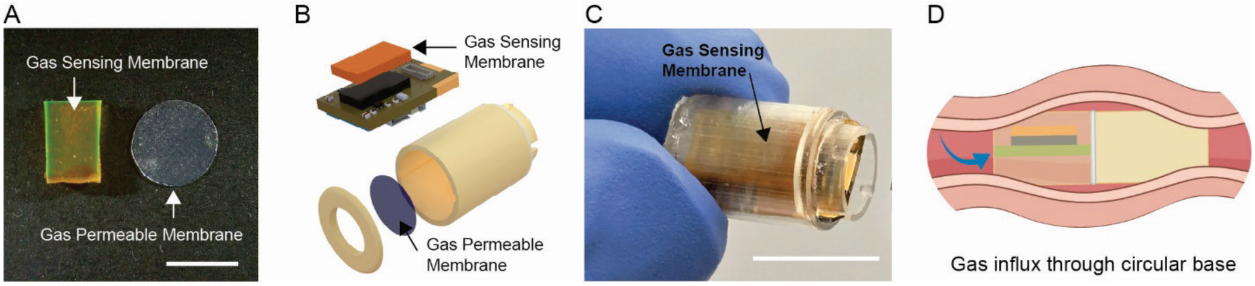

**Fig. S6: Implementation of optoelectronic gas sensing membranes into ESCAPE.** (A) ESCAPE's optoelectronic oxygen gas sensing module requires both a flourescent membrane and a gas permeable membrane (scalebar = 5mm). (B) The gas sensing membrane sits on to of the sensing module, wheras the gas permeable membrane is built into the control capsule segment's cap. (C) Fully assembled control capsule module with both the fluorescent and gas permeable membranes incorporated (scalebar = 10 mm) (D) The gas permeable membrane is placed on the circular base of the capsule to minimize contact with luminal tissues, which can hinder gas diffusion when conformally contacting the membrane. Illustrations in **d** licensed from Adobe Stock under Adobe For Enterprise License

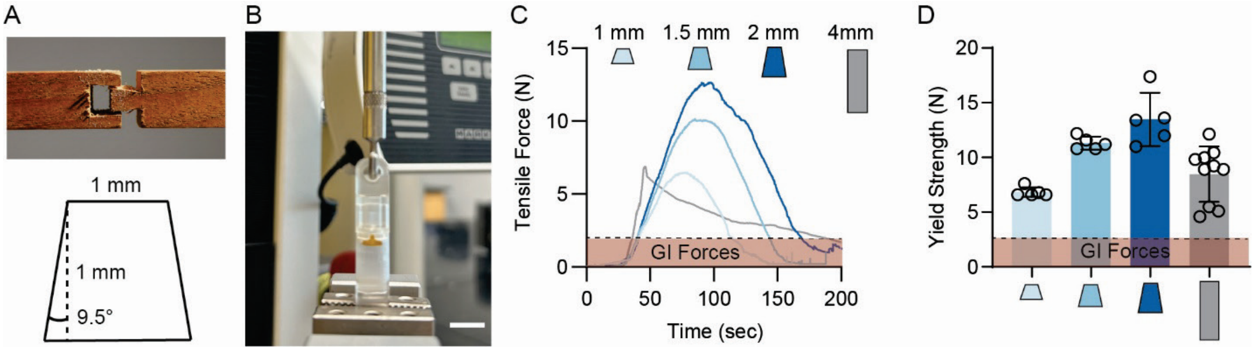

**Fig S7: Design and Characterization of the Dovetail Polymer Interlocks.** (A) The heat severable interlock that joins each capsule segment is inspired by dovetail joinery. Our dovetail channels use a 1:6 ratio commonly used for soft, malleable wood. (B) Capsule seals were tested mechanically using a Mark10 tensile tester (scalebar = 10 mm). (C) Different dovetail lengths were tested at a constant channel top width of 1 mm and compared to a rectangular control channel 4 mm in length. While the rectangular channels exhibit a sharp yield stress when loaded, dovetail channels show significantly increased ultimate tensile strength. A 1 mm dovetail can resist the same tensile force as the 4mm rectangular channel at 1/4 the channel's volume. (n = 5-10 devices, line = Avg) (D) Similarly, dovetail channels exhibit much less variance in yield strength compared to rectangular channels, which may be due to the dovetails ability to engage the tensile modulus of the polymer seal compared to just the shear modulus. A 1 mm dovetail was therefore selected for ESCAPE due to its ability to resist forces experienced in the GI tract (2N) while minimizing the effective volume of the channel (n = 5 - 10 devices, Avg ± SD). Illustrations in **a** licensed from Adobe Stock under Adobe For Enterprise License

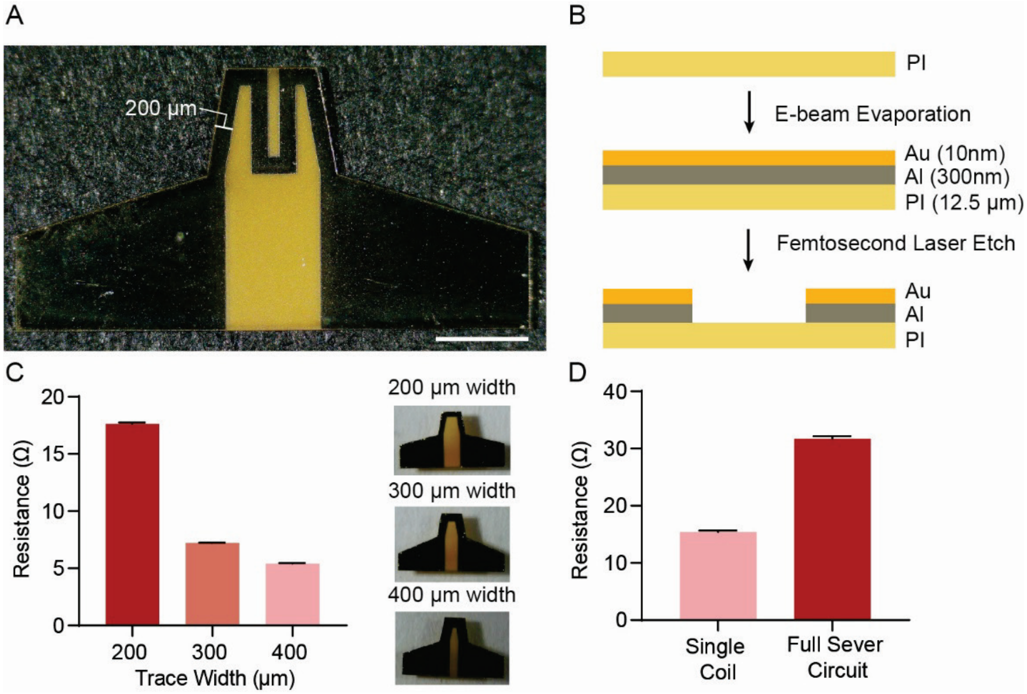

**Fig. S8: Capsule severing uses flexible, thin film, resistive heating coils to generate the necessary heat for actuation.** (A) Design of a serpentine thin film heater (scalebar = 1 mm). Resistance is primarily tuned by adjusting the thickness of the aluminum heating layer. Trace width was fixed at 200 um. (B) Fabrication of thin film heating coils (PI = Polyimide; Au = Gold; Al = Aluminum). (C) Variations in trace width result in resistance variation at the cost of decreased heater surface area (n = 10 coils, Avg ± SD)). Trace width was therefore fixed at 200 um to maximize trace length and effective heater surface area. (D) Thin film heating coils can be inserted with ESCAPE's form factor and soldered to the fPCB to form a two-coil resistive heating circuit when arrayed in series (n = 10 coils, Avg ± SD).

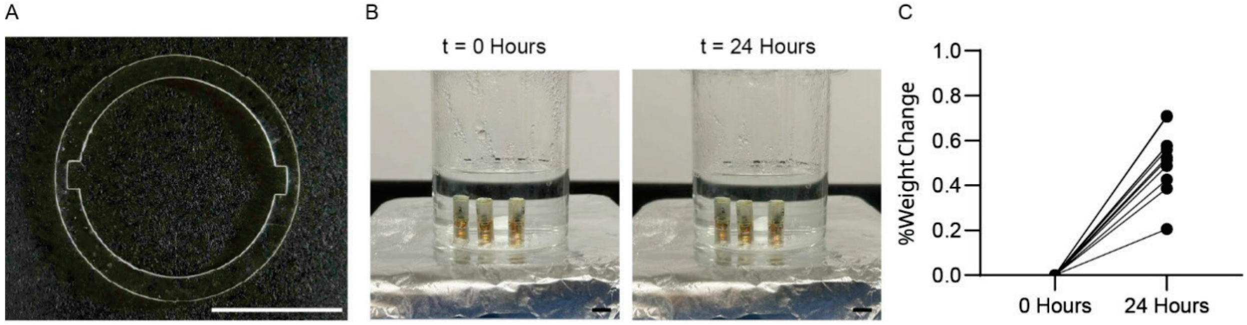

**Fig S9: Watertight PDMS gasket design for ESCAPE, pre-severing. (**A) Design of the watertight gasket (scalebar = 5mm). (B) When submerged in an acidic medium (SGF, pH = 1.3) at 37°C and under a stir rate of 400 rpm, capsules do not exhibit visual indication of leakage of a methylene blue payload over 24 hours (n=10 devices, scalebar = 10 mm). (C) Following the watertightness experiment, capsules exhibited < 1% increase in weight due to moisture influx (n = 10 devices).

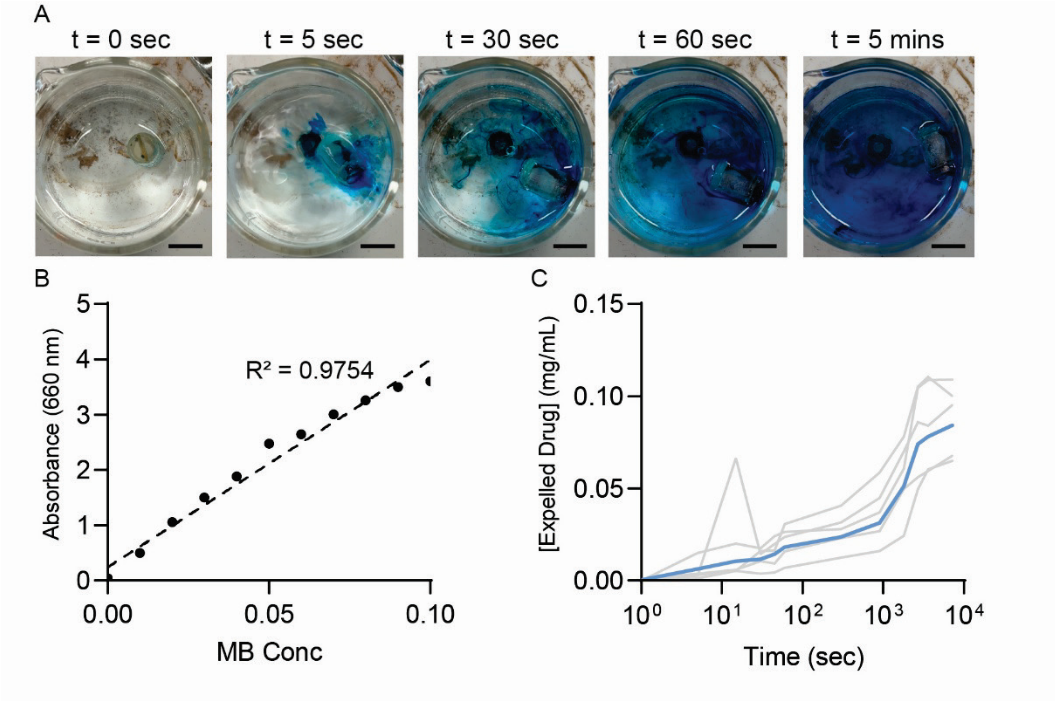

**Fig S10: Proof of concept drug expulsion using Methylene Blue (MB) in DI water.** (A) Visual representation of drug expulsion using a pill containing 5 mg of methylene blue (scalebar = 10 mm). (B) Standard curve of Methylene Blue concentration in DI water (n = 5 curves). (C) Average drug expulsion from ESCAPE over time at 37 ºC (n=5 devices, bold = avg).

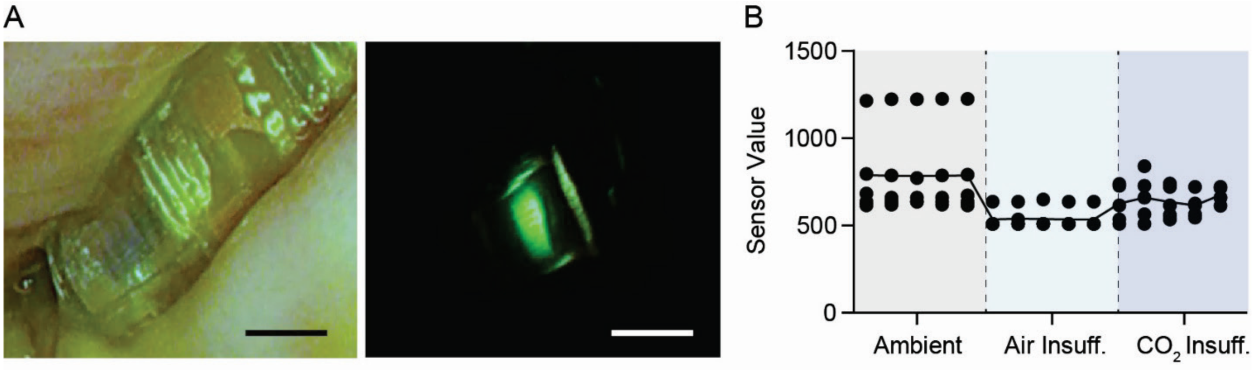

**Fig. S11: In vivo demonstration of optoelectronic sensing.** (A) Representative images of ESCAPE during a sensing cycle (scalebar = 10 mm). Full sensing data collected during ESCAPE deployment (n=5 devices, line = avg). Ambient values collected outside each animal were significantly elevated compared to measurements following air insufflation in the stomach.

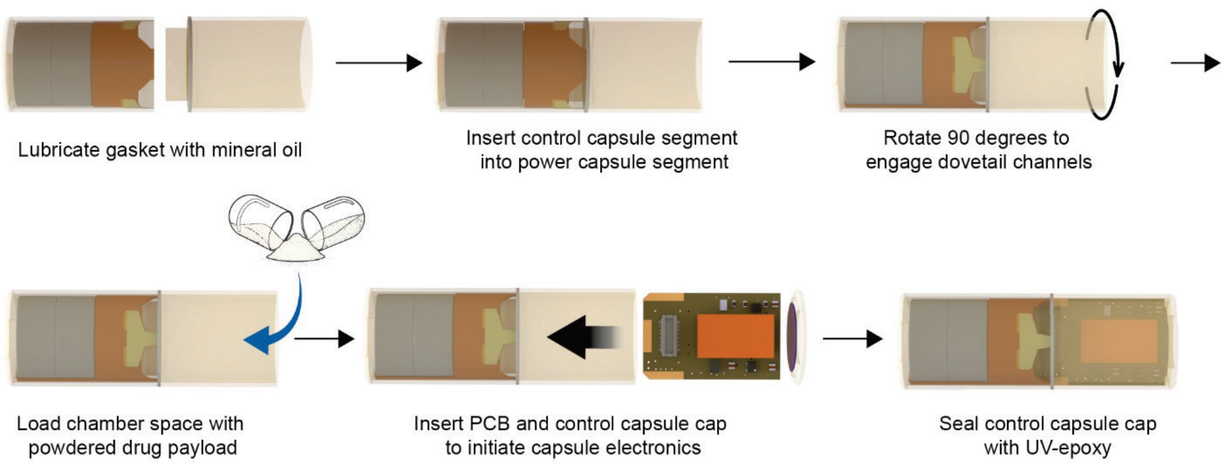

**Fig. S12: Assembly of the ESCAPE platform.** (A) Full capsule assembly can be achieved in 6 stages with simple materials. During assembly, the joined ESCAPE platform can be loaded with powdered drug or any compatible board as determined by the user.

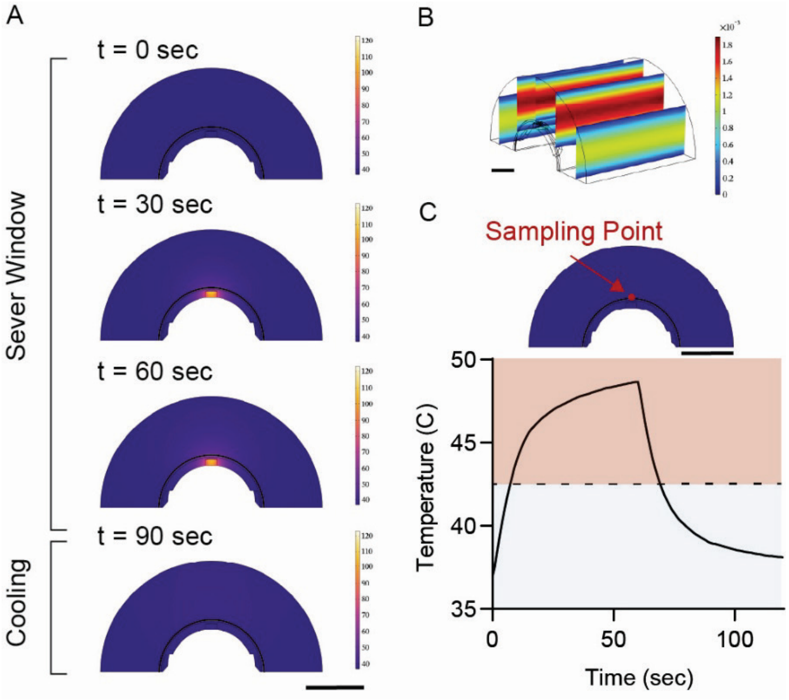

**Fig. S13: COMSOL simulations of ESCAPE heat generation and dissipation in a simulated physiological environment.** (A) ESCAPE's sever circuit produces localized hotspots over a 60 second period. After 60 seconds, heat generation ends and the device cools back to near physiological baseline temperatures after 30 seconds. (B) Laminar flow boundary condition established to mimic chyme flow rate. (C) Temperature sampled from the interface between the capsules outer edge and the aqueous environment over 120 seconds (dashed line = 43º C). Scalebar = 5mm.

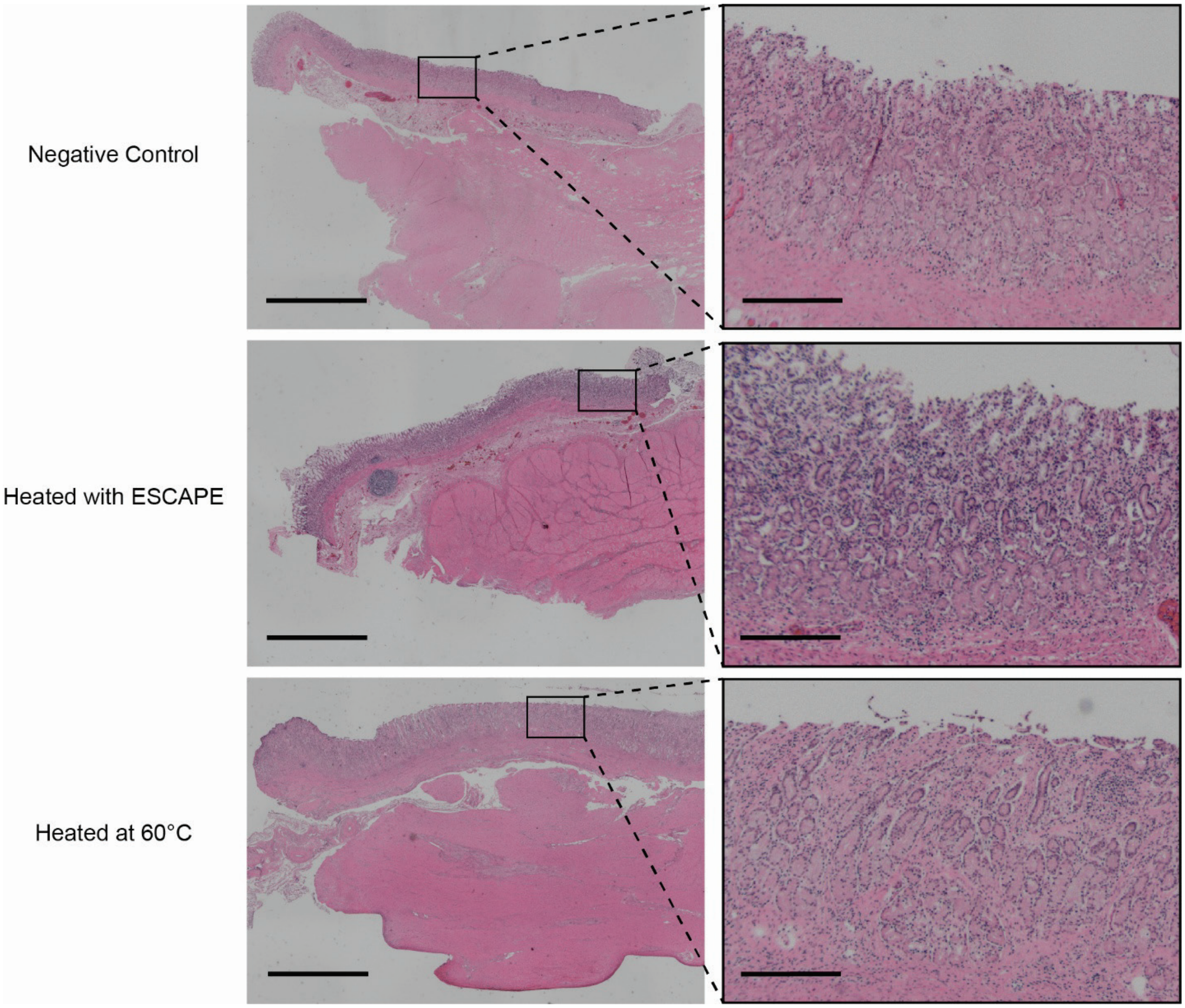

**Fig. S14: ESCAPE thermal dose histology in ex-vivo swine stomach.** (Left) H&E-stained non-heat-treated samples compared to (Middle) samples held in contact with ESCAPE and actuated for 60 seconds and (Right) samples heat at 60°C for 10 minutes. Scalebar = 2 mm (zoomed out); 250 μm (zoomed in)

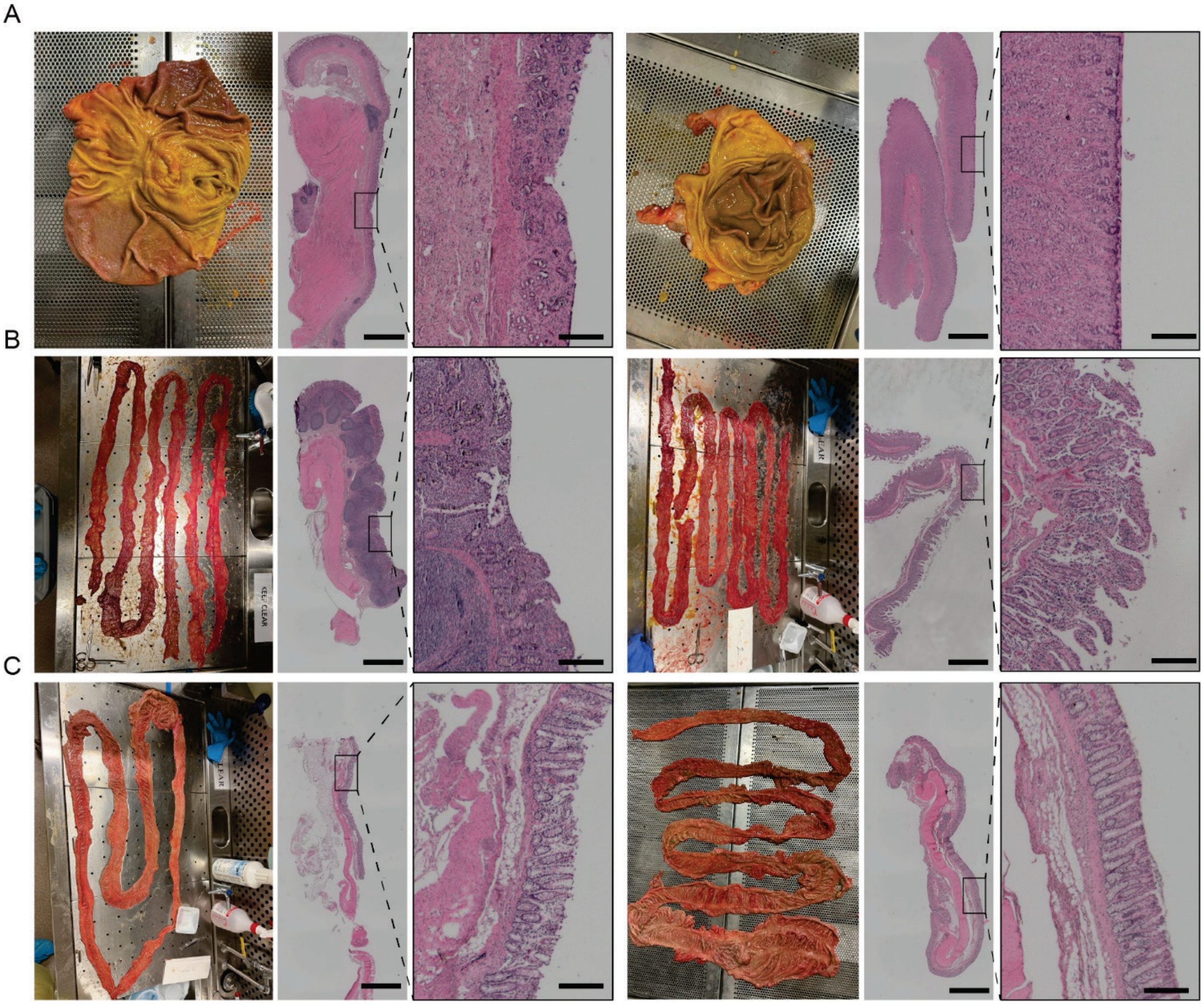

**Fig. S15: Representative images and histology from GI organs following in-vivo evaluation of ESCAPE.** The (A) stomach, (B) small intestine, and (C) colon tissues that ESCAPE passed through in vivo were collected and visualized. No gross damage was observed in any GI organ post-delivery of ESCAPE (n=2 animals). Scalebar = 2 mm (zoomed out); 250 μm (zoomed in)
